# Scalable imaging-based profiling of CRISPR perturbations with protein barcodes

**DOI:** 10.1101/2025.11.11.687767

**Authors:** Krishna Choudhary, Michael T. McManus

## Abstract

Imaging-based CRISPR screens enable high-content functional genomics by capturing phenotypic changes in cells after genetic perturbation. Protein barcodes provide cost-effective, easy-to-implement, and imaging-compatible barcoding for pooled perturbations, yet their scalability has been constrained by the need for arrayed cloning, lentiviral recombination between barcodes and guides, and difficulties in decoding barcodes with high confidence. Here, we introduce poolVis and cellPool, an integrated experimental and computational platform designed to address these limitations. poolVis uses Cre-lox-mediated reconfiguration to position barcode-sgRNA pairs in proximity during viral integration, which greatly reduces barcode shuffling during pooled cloning and delivery. cellPool leverages a scalable computational workflow and the unique aspects of protein barcodes to produce unpooled image galleries from multi-terabyte scale datasets. Applying this platform to single- and double-CRISPRi profiling of cell-cycle genes and chromokinesins in the MCF10A cells uncovered established and previously unrecognized phenotypes, including nuclear morphology changes and reciprocal sign epistasis in DNA damage.

## INTRODUCTION

Microscopic imaging provides rich information in space and time, revealing phenotypes that DNA sequencing cannot capture. Imaging-based functional screens have propelled cell biology and drug discovery.^1,2^ Recent confluence of CRISPR technologies with high-throughput, multiplexed imaging has renewed interest in pooled imaging-based profiling of genetic perturbations.^2^

Profiling thousands of perturbations in isolation is labor-intensive. Optical barcoding reduces labor by pooling perturbations. To be practical, barcodes must be robustly decodable with controlled error rates. Current approaches rely on either RNA or protein barcodes.^2^

While RNA barcoding via *in situ* sequencing offers valuable spatial information, it is costly and requires multiple amplification steps, high-temperature incubations, artisan in situ sequencing techniques, and potentially cell-line-specific spot detection methods.^3,4^ Protein barcodes, in contrast, use immunostained epitope tags fused to stable proteins expressed by strong promoters, and are compatible with available automation solutions.^5,6^ Their nuclear protein (e.g., H2B) fusions give bright, easy-to-detect signals even at low magnification. However, an unexpected and often underestimated burden is antibody panel optimization; failing this, debarcoding performance may suffer.

Existing lentiviral vector designs position protein barcodes and sgRNAs far apart (1 kb or more) on the backbone and are prone to recombination during viral packaging, a major source of barcode-sgRNA mispairing. In addition, libraries with identical coding sequence for an epitope tag across barcodes can exacerbate lentiviral recombination and PCR-mediated template switching during amplification, further compounding linkage errors^7–10^. Consequently, the scalability of pooled protein-barcoded libraries has remained limited.^5,11^ Although one large-scale protein barcode study has been reported^6^, it did not involve CRISPR screens, where the absence of gene-specific phenotypes simplified interpretation despite potential barcode shuffling. Thus, the need to scalably generate protein-barcoded libraries, read barcodes cost-effectively, and ensure performance quality despite staining variation remains unmet.

To address these challenges, we present a platform addressing these needs. It combines a wet-lab component, poolVis (Fig. 1a), with a computational pipeline, cellPool (Fig. 2a). Our platform differs from prior art by (i) use of Cre-lox reconfiguration that maintains ~50 bp protein barcode-sgRNA proximity in poolVis vectors to suppress template switching and barcode shuffling,^7,12,13^ (ii) a TATAlox-U6 architecture and non-homologous barcode coding sequences to stabilize barcode and sgRNA expression with Cre induction in the transduced cells, and (iii) cellPool, a Nextflow-based workflow that performs high-quality image processing and multivariate model-based debarcoding to output unpooled single-cell galleries for quality control and interpretation.

**Figure 1.**
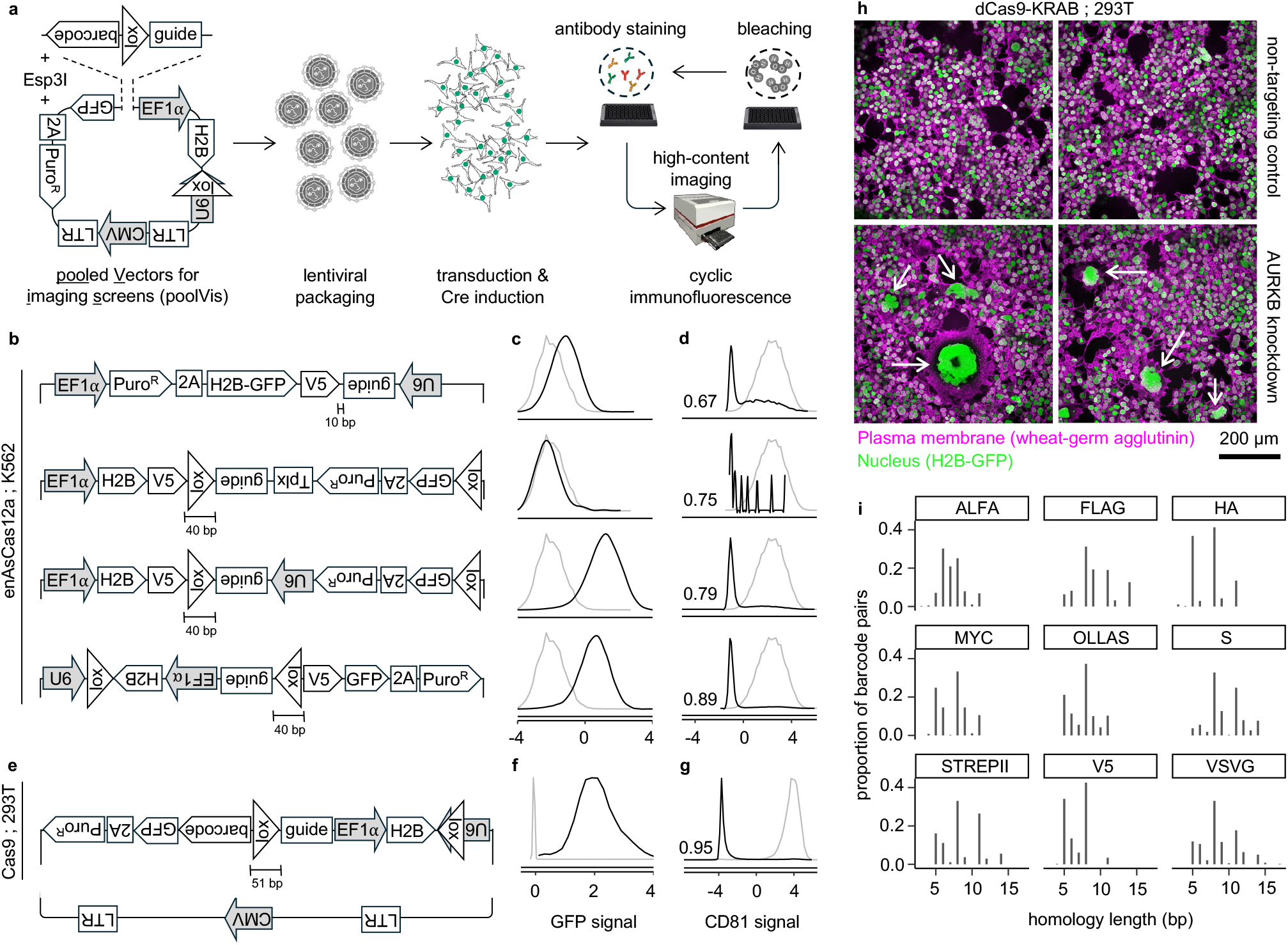
poolVis is a Cre inducible CRISPR platform that minimizes the randomization of protein barcode-guide RNA pairs due to recombination in pooled lentiviral libraries. **a**, Overview of the poolVis workflow. **b, e** Lentiviral vector designs for Cas12a- and Cas9-based screens. **c, f**, GFP signal from the design positioned in the same row to the left, scaled and centered with respect to the background green fluorescence in untransduced control cells (gray). **d, g**, *CD81* signal from the design positioned in the same row to the left, scaled and centered with respect to the non-targeting control (gray). **h**, Imaging of cells with *AURKB* knockdown shows enlarged nuclei (examples highlighted with white arrows). **i**, Distribution of pairwise homology lengths in the set of coding sequences designed for each epitope tag in the barcode library. Short homology tracts indicate minimal shared nucleotide identity between barcodes, confirming effective homology reduction by codon optimization.

**Figure 2.**
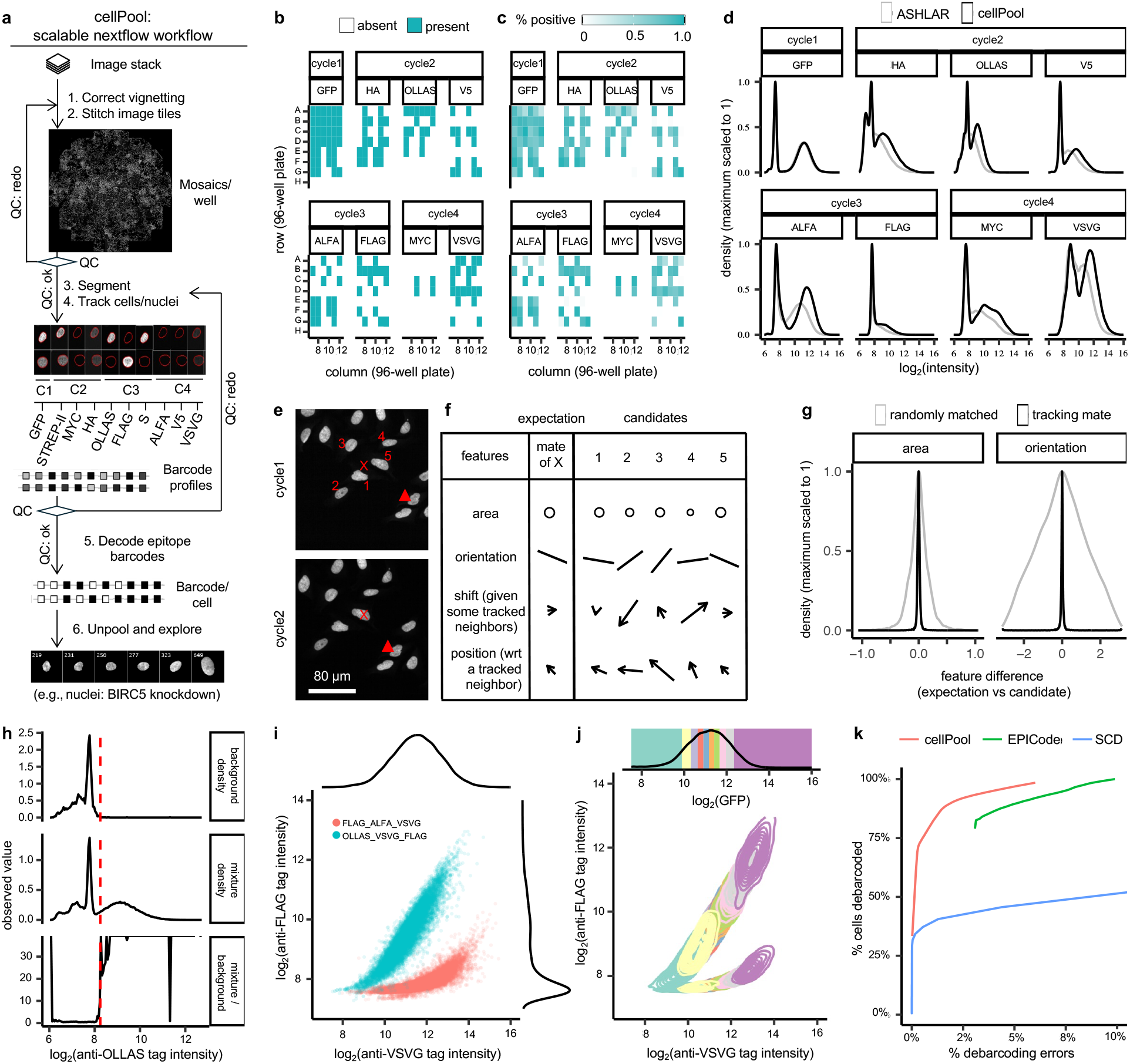
cellPool enables scalable computational analysis to reverse the pooling of CRISPR perturbations. **a**, Overview of the cellPool workflow. **b**, Plate map showing the presence or absence of each barcode element (GFP and epitope tags) in 19 barcoded U2OS samples for benchmarking cellPool (one barcode per well) and untransduced U2OS cells. **c**, cellPool debarcodes the samples with high accuracy. The colormaps are showing percentages of cells positive for each barcode element in a well. **d**, The signal-background mixture distributions estimated by ASHLAR (gray) and cellPool (black). **e-f**, cellPool tracks cells from later imaging cycles to find their “mate” cell in the previous cycles. The tracking is initialized with image coordinates after registration of image mosaics, e.g., the mate of a nucleus at position labeled with red X in cycle 2 is searched for in the cycle1 image starting from the same position with respect to the top left corner. The number of nuclei that are candidates for matching is user-defined (labeled with red numbers in cycle1 image). The nuclei with high-confidence tracking mates (example tagged with red triangle) available increase in number with every iteration of tracking. **e-g**, Clonal expansion of adherent cells leads to neighboring cells being more likely to have similar area, but orientation shows higher randomness. **h**, cellPool initializes debarcoding with univariate analysis of each epitope tag by comparing the signal-background mixture distribution with the background only distribution. **i**, The immunostaining signal for an epitope tag is dependent on other epitope tags included in the protein barcode. Example, the FLAG tag signal is much weaker in one of the shown barcodes. **j**, cellPool models the barcode profiling data as derived from a mixture of multivariate GFP-dependent skew-elliptical distributions. **k**, cellPool outperforms other existing methods for debarcoding the protein barcodes.

We apply poolVis and cellPool to profile 336 single and 2,016 double CRISPR interference (CRISPRi) perturbations. To minimize antibody use and imaging time, we arrayed minipools of 84 barcoded sgRNAs, readable in three cycles. This design scales to >1,000 sgRNAs in a dozen wells, focused on specific pathways, and by extrapolation to genome-scale libraries using two 96-well plates. We recover known and previously unrecognized phenotypes, validating hits individually. Together, these results establish a cost-effective and scalable platform for imaging-based profiling of CRISPR perturbations.

## RESULTS

### poolVis enables pooled CRISPR perturbations with protein barcodes

We developed poolVis to bring pooled barcode-sgRNA pairs closer than 100 bp. Our prototype considered Cas12a-CRISPR knockout because of shorter Cas12a scaffold than Cas9.^14^ Here, barcodes were fused to EF1α-driven H2B-GFP, while the human U6 (hU6) promoter downstream of stop codon drove the sgRNA (Fig. 1b, top design). In flow cytometry with enAsCas12a-expressing K562s and CD81-targeting sgRNA, GFP signal and CD81 knockout were poor (Fig. 1c-d top row; Supplementary Table 1a-b; Supplementary Fig. 1a), suggesting EF1α and U6 transcriptional interference.^15^

To resolve this, we leveraged Cre-lox reconfiguration. We added lox71 between barcode and sgRNA in the oligonucleotide inserts and lox66 in the vector backbone. After viral integration, Cre inverts the floxed cassette, rearranging the barcode and sgRNA, and lox66/71 unidirectionally converts to functionally inert loxP/72. This preserves sgRNA-barcode proximity until Cre induction, after which transcriptional interference can be prevented.

To maximize barcode and sgRNA expression, we evaluated several constructs. A one-transcript design, where Cre recombination produced a single mRNA encoding the H2B-GFP barcode and the sgRNA separated by a stabilizing Triplex motif (Fig. 1b, second design), was inspired by prior mRNA-embedded guide strategies.^16^ However, puromycin selection revealed poor viability, suggesting mRNA destabilization (Fig. 1c-d, second row). We therefore developed designs that reconfigure barcodes and sgRNAs into separate transcripts.

Building on these results, we tested configurations where Cre recombination repositions U6 either downstream or upstream of EF1α (Fig. 1b, third and fourth designs). Using *CD81*-targeting sgRNA, we observed the best combined CRISPR activity and barcode signal with U6 upstream of EF1α (Fig. 1c-d, bottom rows). Here, 40 bp barcode-sgRNA separation reconfigured into highly functional units.

To generalize the design, we next adapted it for Cas9 and dCas9-KRAB systems. The recombined loxP sequence is fused to the Cas12a-sgRNA in the best design above and removed by Cas12a, which Cas9 cannot do. Also, the long Cas9 scaffold should be in the vector backbone. We therefore implemented a TATAlox-mU6 promoter, embedding loxP downstream of the mouse U6 (mU6) proximal element.^17^ To promote unidirectional recombination, we engineered TATAlox66/71 variants (Supplementary Table 1c). To improve lentiviral titer and eliminate significant recombination, we oriented EF1α tandem with CMV and reduced barcode-sgRNA separation to 51 bp (Fig. 1e). In HEK293Ts, these design implementations yielded efficient Cas9 *CD81* knockout, dCas9-KRAB *CD81* repression, and barcode signals (Fig. 1f-g; Supplementary Fig. 1b-c), and *AURKB* repression caused expected nuclear defects (Fig. 1h). To further improve gene knockdown, we additionally produced TATAlox-hU6 promoters (Supplementary Fig. 1d-f; Supplementary Table 1d). These results establish poolVis for pooled protein-barcoded Cas12a, Cas9, and dCas9-KRAB libraries.

### Design of non-homologous coding sequences for protein barcodes

Template switching and barcode recombination can occur between identical nucleic acid sequences during lentiviral reverse transcription or PCR amplification. To reduce these artifacts, we codon-optimized epitope tag and barcode sequences to minimize homology. A brute-force pairwise comparison to minimize library-scale sequence homology is computationally impractical, e.g., V5-tag admits >10^8^ possible coding sequences. We therefore designed a heuristic algorithm to generate 84 minimally homologous barcode coding sequences. The algorithm fragments each peptide into short motifs, enumerates codons per motif, and iteratively assembles coding sequences 5′→3′ while minimizing homology to previously sampled coding sequences. This process produced a diverse set of non-homologous barcode coding nucleic sequences resistant to template switching and recombination (Fig. 1i, Supplementary Table 1e).

### cellPool enables scalable analysis of high-content imaging data

cellPool is a Nextflow^18^ pipeline that transforms multiplexed imaging data into unpooled image galleries of debarcoded cells (Fig. 2a). Starting with illumination correction, imaging tiles are stitched together, nuclei are segmented, tracked across cycles, barcode intensities quantified, and debarcoded, and single-cell galleries are generated. Containerization ensures reproducibility and straightforward deployment on high-performance computing systems.

To benchmark cellPool, we transduced 19 barcodes individually in U2OS in a pre-determined plate map (Fig. 2b). We imaged and analyzed the samples without the map. cellPool recovered barcode identities with high sensitivity and specificity (Fig. 2c) attributable to our mosaicking, tracking, and debarcoding methods.

### cellPool yields high-quality integration of iterative imaging data

ASHLAR is a state-of-the-art method for multiplexed imaging integration using phase correlation,^19^ which estimates translational misalignments but not other transformations. It has been validated for five microscopes, predominantly pathology slide scanners. Our experiments used a well-known high-content microscope where ASHLAR frequently misaligned images (Supplementary Video 1), increasing signal-to-background overlap (Fig. 2d). Hence, cellPool stitches image tiles from each cycle, but skips pixel-to-pixel registration across cycles. This ensures robustness to instrumental variations. cellPool mosaicking is based on ASHLAR algorithm and outputs detailed quality reports and videos showing the alignments to enable human-in-the-loop fine tuning (Supplementary Fig. 2-3; Supplementary Videos 2 and 4).

cellPool unpools registered images, but internally skips pixel-to-pixel registration. This ensures robustness to unforeseen artifacts. cellPool integrates the single-cell information across imaging cycles by tracking in multiple iterations. It leverages invariance of fixed-cell nuclear orientations in the first tracking iteration (Fig. 2e-g). Although largely random, neighboring nuclei in dividing or nascently divided cells may present identical orientations causing tracking errors, which are identified based on arrangement of neighboring nuclei and corrected by positioning relative to the nuclei correctly tracked in the previous iteration. Our approach gives near perfect tracking confirmed by 100s of randomly sampled tracking mates (Supplementary Videos 3 and 5).

### cellPool enables accurate decoding of protein barcodes

cellPool uses model-based clustering for debarcoding premised on the following observations. First, artifacts such as antibody aggregates may lead to multimodality, and poor signal may cause unimodality and heavy tails (Fig. 2h). Second, the multivariate barcode distributions are not elliptical, symmetric, or convex (Fig. 2i). Third, the epitope intensities depend on H2B-GFP signal and have covariance patterns that can be leveraged for debarcoding (Fig. 2j).

Debarcoding starts with a high-stringency non-parametric univariate thresholding by comparing signal-background mixture with background only populations (Fig. 2h). This reduces the downstream computational complexity by partial debarcoding without invoking distributional assumptions. Next, we group the nuclei into 5-10 sets by H2B-GFP intensity (Fig. 2i-j) and learn covariance patterns by expectation-maximization (Supplementary Fig. 4), which extends prior work on skew-elliptical distributions for flow cytometry data^20^ to protein barcodes. This gives a barcode score matrix and assignments per nucleus. In benchmarking test, cellPool consistently outperformed EPICode^6^ and single-cell deconvolution (SCD)^21^ (Fig. 2k) as expected given explicit covariance modeling.

### CRISPRi screen of cell-cycle genes in MCF10A

Cell cycle regulation ensures controlled growth and DNA replication. MCF10A breast epithelial cells are a physiologically relevant model to study this process given their stable near-diploid karyotype and intact cell-cycle checkpoints with wild-type *TP53* and *RB1*.^22^ Profiling cell-cycle perturbations in MCF10As can reveal distinct functions compared to tumorigenic HeLa or A549.^23^

We applied poolVis and cellPool to profile 336 CRISPRi perturbations of cell-cycle targets (Fig. 3a; Supplementary Table 1f-g). Guides were arranged as four minipools of 84 barcoded sgRNAs each. MCF10As with genomically integrated dCas9-KRAB and ERT2-Cre-ERT2 were processed as in Fig. 1a. We used a TATAlox-mU6-driven sgRNA cassette here, which achieved 71% CD81 knockdown relative to 98% under hU6 (Supplementary Fig. 5a), indicating partial loss-of-function for majority targets as expected for the mU6 promoter.^24^

**Figure 3.**
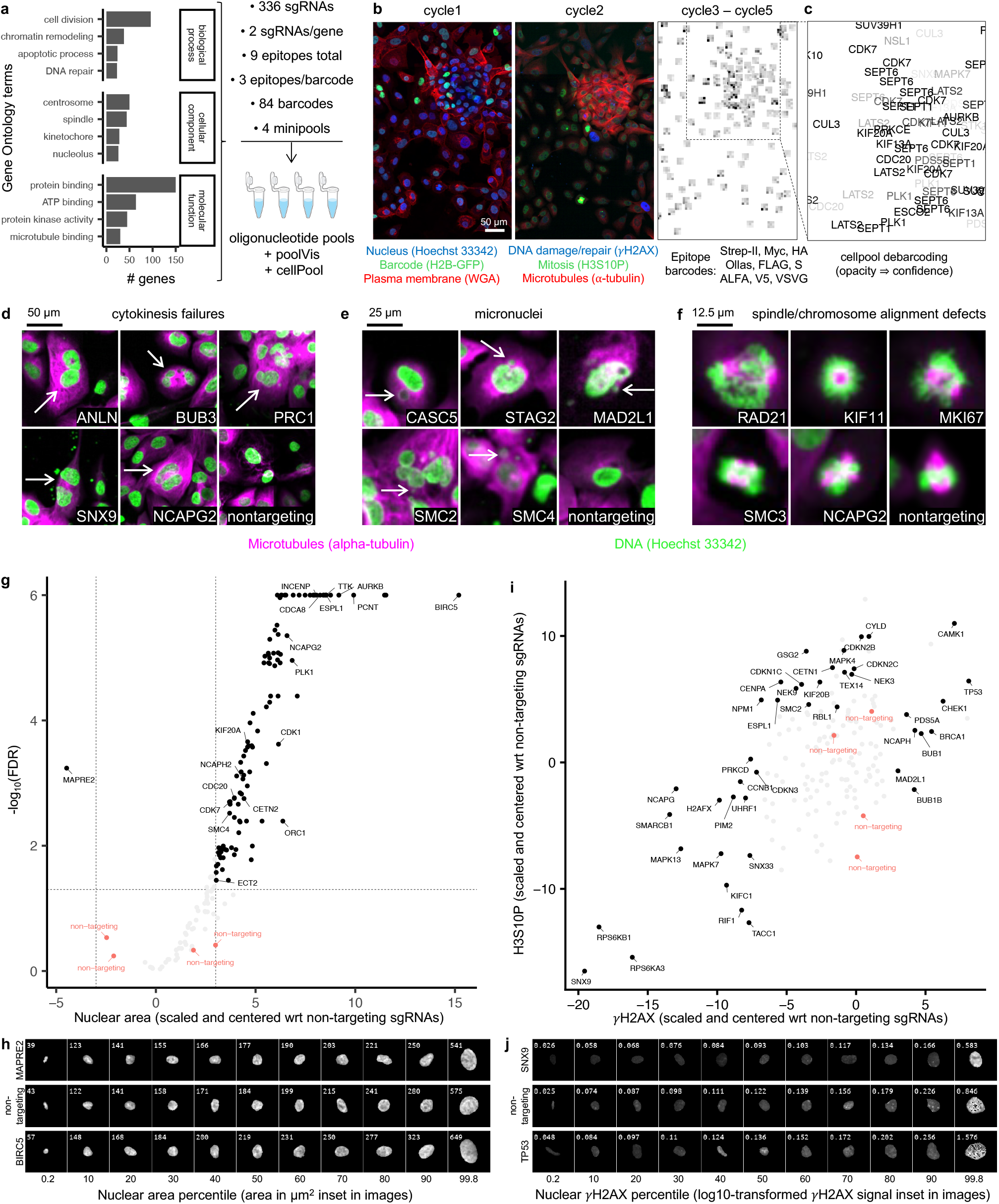
poolVis and cellPool enable screen of cell-cycle regulating genes with multiplexed imaging readout. **a**, Gene ontology terms associated with the targeted genes and breakdown of the minipools. **b**, Multiplexed imaging panel showing pooled perturbations. The barcode profiles are shown as squares with nine tiles with each tile showing the median fluorescence intensity per nucleus for the epitope tags. **c**, cellPool outputs the identities of genetic targets per nucleus. **d-f**, Representative phenotypic effects of CRISPR knockdowns unpooled by cellPool. **g**, Volcano plot showing the perturbations that impact the nuclear area. **h**, Representative images of nuclei with perturbations that increase or decrease the nuclear area. **i**, Scatter plot comparing the shift in H3S10P and γH2AX signal across genetic targets. **j**, Representative images of nuclei with perturbations that increase or decrease the nuclear γH2AX signal.

We profiled nuclei, plasma membrane, and H2B-GFP, followed by DNA damage/repair using anti-γH2AX, mitotic index using anti-phospho-histone H3 (H3S10P), and microtubules using anti-α-tubulin (Fig. 3b-c). Then, barcode profiling took three iterations, costing approximately 10-fold less per cell compared to RNA barcodes (Supplementary Table 1h). Analysis reports confirmed high-quality image processing (Supplementary Videos 4-5). With debarcoding error rate estimated at 1.4% for our confidence threshold (Fig. 2k), 454,943 debarcoded nuclei were considered for downstream analysis, excluding uninfected and multiply infected cells (median: 996; Supplementary Fig. 5b). Unpooled images revealed expected phenotypes (Fig. 3d-f).

Next, we conducted differential analysis (Supplementary Table 2). As expected for cell-cycle genes, most knockdowns caused enlarged nuclei relative to the non-targeting controls likely due to checkpoint activation and/or endoreplication^25^ (Fig. 3g). The chromosomal passenger complex genes, *BIRC5, AURKB, CDCA8*, and *INCENP*, were among the top hits (Fig. 3g-h). *MAPRE2* was the only hit for smaller nuclei, which is consistent with a recent RNAi screen.^26^

The screen revealed established hits for DNA damage/repair (Fig. 3i). *TP53* knockdown increased the integrated nuclear γH2AX signal the most (Fig. 3i-j). Other established hits for increased DNA damage were *CHEK1* and *BRCA1* (DNA repair), *NCAPH* (condensin I), *PDS5A* (cohesin regulator), and *BUB1, BUB1B* and *MAD2L1* (spindle assembly checkpoint). As expected, *H2AX* knockdown reduced γH2AX signal. In general, cell-cycle dysregulation increases DNA damage due to mechanisms such as replication stress, impaired DNA repair, checkpoint loss, and cytokinesis failures. However, γH2AX staining measures both DNA damage and repair, specifically, functional expression, trafficking and assembly of γH2AX at the damage sites. For multifunctional genes, γH2AX signal summarizes multiple processes.

Interestingly, more perturbations reduced γH2AX signal than enhanced. Though *UHRF1* maintains genome integrity, its knockdown reduces γH2AX assembly at damage sites^27^ explaining the reduced γH2AX signal. Since microtubules help transport DNA repair proteins,^28^ microtubule-associated hits such as *KIFC1, TACC1*, and *MAPRE2* are expected. *RPS6KA3* and *RPS6KB1* phosphorylate ribosomal protein S6 to stimulate translation. Their knockdown causes overall reduced cellular protein, plausibly including γH2AX. *NPM1*, involved in ribosome biogenesis, also facilitates DNA repair. Reduced γH2AX after *NPM1* knockdown suggests that its partial loss-of-function impacts translation more than DNA repair. Similarly, multifunctional genes *SNX9* and *SNX33*, well-known for roles in cytokinesis, reduced γH2AX (Fig. 3j). Overall, our data highlight that for multifunctional genes, perturbation dose characterization facilitates interpretations. Alternatively, more repair proteins could be profiled with our platform.

H3S10P peaks during the late G2/M phase. Perturbations that arrest cells in the late G2/M phases should increase the H3S10P signal. Since only a minority of cells is expected in the late G2/M phases, there was higher variability in mean H3S10P of nontargeting controls than for mean γH2AX (Fig. 3i). Despite this, we uncovered established hits, e.g., *GSG2* and *CENPA*, and observed correlated mean H3S10P and γH2AX. Further, we identified mitotic nuclei based on H3S10P thresholding and computed the mitotic indices (Supplementary Fig. 5c). Given good coverage, the mitotic indices for the two guides targeting the same gene were correlated (Supplementary Fig. 5d), and in the expected range (1.02%) for the non-targeting and untransduced cells. *BRCA2, MAPK7, PRKCE, RB1*, which regulate cell-cycle progression and mitotic entry, and *KIFC1*, which regulates spindle organization and mitotic progression, were the top hits for lowered mitotic index (FDR <0.1). No hits increased mitotic index. Similar analysis revealed *KIF2A*, a microtubule depolymerase, as the top hit for enhanced α-tubulin staining (Supplementary Table 2). Overall, these observations support that our data is high-quality.

To validate, we cloned sgRNAs targeting selected genes and nontargeting sgRNAs individually in a control vector under hU6. Then, fully arrayed samples were processed to image cycles 1-2 as in Fig. 3b and analyzed by cellPool. For confident hits identified by FDR-adjusted p-values and single-cell coverages, the arrayed minipools and fully arrayed screens largely agreed (Supplementary Fig. 6), confirming that poolVis and cellPool deliver high-quality imaging-based profiles and robust analysis.

### Arrayed minipools enable low-cost double-CRISPRi screens

Chromokinesins are microtubule-based motors that ensure proper chromosome segregation.^29^ Their functional epistatic interactions are of significant interest, but have not been profiled at scale by imaging. To investigate epistasis, we extended poolVis for dual-sgRNA expression by adding another U6 promoter (Fig. 4a) and generated six backbones, each containing an hU6-driven spacer targeting both P1/P2 isoforms of *KIF15, KIF4A*, and *ATM*, or *KIF22* isoforms individually, or a non-targeting control. We validated each spacer individually (Supplementary Fig. 7). The four minipools in Fig. 3a were inserted for a library of 2,016 double perturbations (Fig. 4b-c). Then, we transduced MCF10A cells, induced Cre, grew cultures, and imaged four fixed plates according to Fig. 3b. cellPool debarcoded 2,159,003 nuclei (median: 751; Fig. 4d; estimated error rate: 1.4%). Technical replicates agreed closely for normalized γH2AX, H3S10P, and α-tubulin, and *H2AX* knockdown robustly reduced γH2AX (Supplementary Fig. 8).

**Figure 4.**
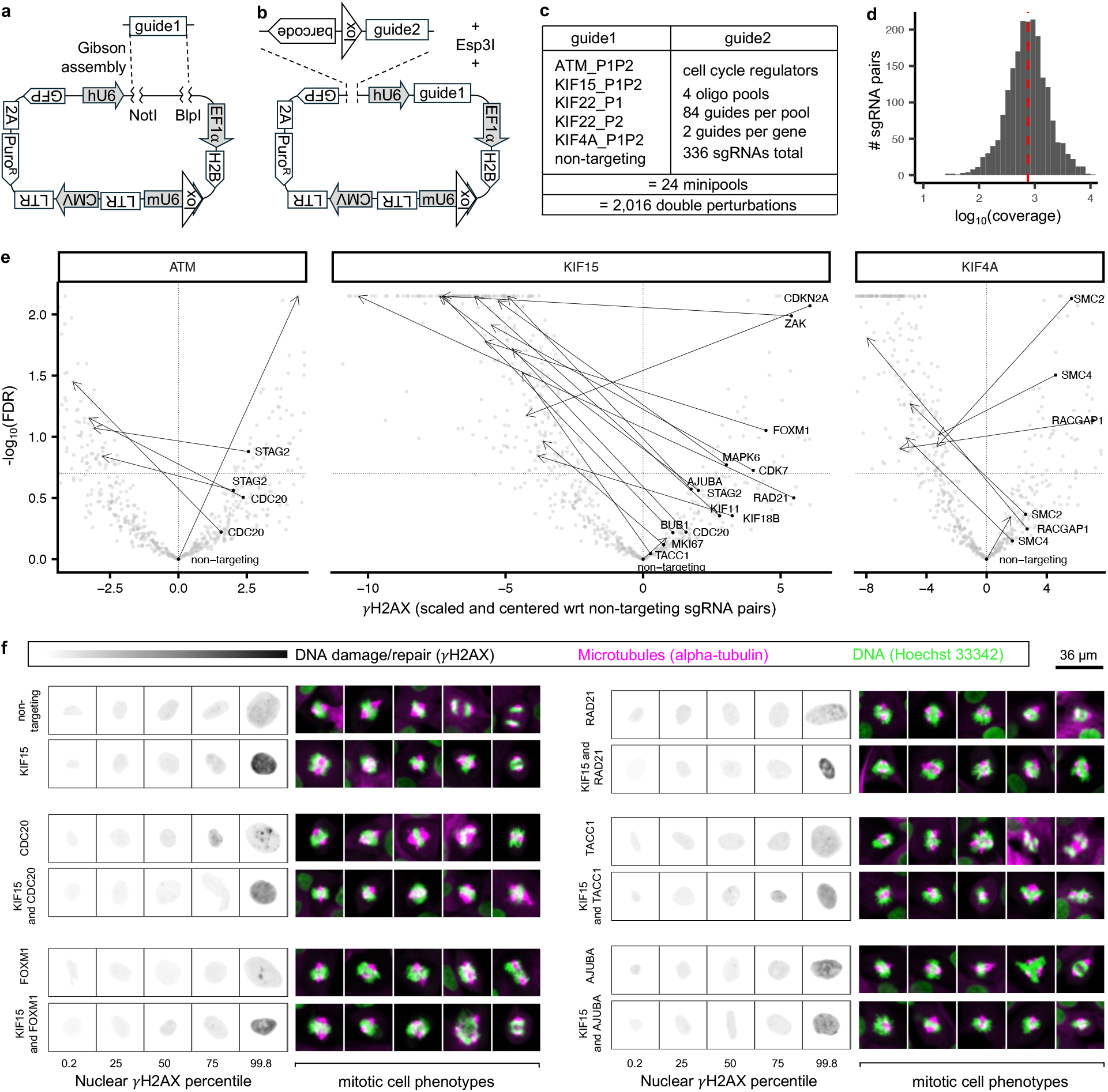
poolVis and cellPool reveal epistatic interactions underlying DNA damage and repair in the MCF10A cells. **a-b**, Cloning design for a lentiviral vector backbone carrying a single CRISPR guide sequence and insertion of pooled guides at the poolVis library entry site. **c**, Breakdown of the 2,016 combinatorial perturbations in the current screen. **d**, Coverage distribution of the guide RNA combinations. Red dotted line is the median. **e**, Volcano plots with annotations highlighting epistatic interaction hits (gene target labels at the base of the arrows). Each arrow connects the point representing the gene target when inserted in the backbone with nontargeting guide1 (arrow base) to the point representing the same gene target in the backbone with a targeting guide1 (arrow head). For *ATM* and *KIF4A*, both guide2 spacers targeting the same interactor are shown. For *KIF15*, single guide2 spacer is shown for each interactor. **f**, Representative images of DNA damage patterns and mitotic nuclei for the epistatic interaction hits.

We computed epistasis scores as differences of double- and single-knockdown effects, and a null distribution from bootstrapping non-targeting pairs.^30^ The scores skewed negative relative to the null (Supplementary Fig. 9a). Since apoptosis is triggered if DNA damage accumulates beyond a threshold given the wild-type *TP53* in MCF10As, phenotypic states with DNA damage equaling the addition of single-knockdown effects may not be accessible to the perturbed cells. This means that typical methods for fitness-based epistasis analysis may not apply to DNA damage data, and new methods for imaging-based phenotypes are under active development.^2^ Deep learning-based methods do not yet outperform simple linear baselines for such analysis.^31^ Hence, to evaluate the biological information quality in our data, we called high-confidence epistatic interactions if double-knockdown effects were significantly negative for both guides (FDR<0.2) while single-knockdown effects were positive (known as reciprocal-sign epistasis^32^ or “opposite” interactions^31^; Fig. 4e). This is a simple, biologically interpretable criterion that avoids assumptions based on prior fitness-based studies while highlighting reproducible epistatic interactions.

For *KIF4A*, interactions identified with *SMC2* and *SMC4* (condensin I subunits) and *RACGAP1* agree with roles in chromosome-axis organization^33,34^ and central-spindle function,^35^ respectively. We often observed cytokinesis defects for *RACGAP1* knockdown; however, manual annotation of 4,135 single-cells suggested that multinucleation alone did not explain *KIF4A-RACGAP1* interaction (Supplementary Fig. 9b-c; Additional Data 1). *ATM* interactions with *STAG2* and *CDC20* were also recovered, consistent with checkpoint and repair pathway functions.^36,37^ *KIF15* showed the most hits, including *MKI67, BUB1, CDC20*, and *KIF18B*, supported by public data^38^ (Supplementary Table 3). Other predicted *KIF15*-interactors were *RAD21* and *STAG2* (cohesin), *MAPK6* and *ZAK* (MAPK superfamily), *AJUBA* (spindle assembly checkpoint), *TACC1* (microtubule stabilizer), *CDK7* and *CDKN2A* (CDK regulators). *KIF15* and *KIF11* are partially redundant; their simultaneous loss-of-function shows monopolar spindles. One *KIF11* spacer narrowly missed our high-confidence threshold despite negative epistasis score, reflecting partial repression (Supplementary Table 4). No epistasis was identified for *KIF22* isoforms, suggesting mutual compensation. Single-cell galleries revealed extreme damage phenotypes even when mean γH2AX decreased, underscoring the value of unpooled visualization (Fig. 4f).

A fully arrayed experiment validated the *ATM-* and *KIF15*-centered epistasis hits where *H2AX* targeting controls showed strong γH2AX reduction indicating robust CRISPRi performance, and the epistasis scores were again strongly negative relative to the null (Supplementary Fig. 9d-e). A representative set of mitotic nuclei from the arrayed plates is shown in Fig. 4f. Overall, these results establish our platform for scalable imaging-based profiling of double perturbations.

### Additional resources for screens with poolVis and cellPool

To facilitate adoption, we generated vectors for wt-Cas9, wt-Cas12a, and dCas9-KRAB engineering (Supplementary Table 1a). Common Cas vectors carry HA tags, conflicting with HA-based barcodes. Users may swap out the HA-tag in barcodes, or use our lentiviral/piggyBac vectors lacking HA or carrying alternative tags for Cas engineering. piggyBac vectors also enable one-step engineering of Cas and ERT2-Cre-ERT2 enzymes.

## DISCUSSION

Protein barcodes enable scalable imaging-based profiling of genetic perturbations, yielding spatially resolved information inaccessible by sequencing. Previously, they were limited by low-throughput cloning and poor debarcoding. poolVis and cellPool address these limitations.

poolVis leverages a Cre-inducible design, a defining feature that enables temporal control of barcode and guide expression while maintaining a compact vector design. This feature will be valuable for studies requiring controlled single- or sequential double-perturbations. In addition, poolVis separates spacers and barcodes by ~50 bp during lentiviral packaging, effectively preventing recombination that complicates barcode-based pooled screening.^7,12,13^ Our barcodes are read in three imaging iterations. More iterations can profile proteins, DNA, or RNA. The present library contains 84 barcodes, scalable further with more epitopes/backbones, although eventually imaging throughput becomes limiting. While in situ sequencing costs are expected to decline, immunostaining-based protein barcode detection currently reduces the per-cell reagent cost to roughly one-tenth that of RNA barcodes, and enables broader experimental replication given the specialized nature of in situ sequencing workflows.

cellPool automates the analysis of multi-terabyte high-content imaging data. It includes detailed quality control and human-in-the-loop checkpoints to verify the intermediate results. Leveraging Nextflow and Docker supports reproducible analysis on high-performance computing platforms common in imaging cores. The pipeline incorporates robust mosaicking and across-cycle integration methods and a multivariate debarcoding model tailored to the statistical properties of protein barcodes. By presenting unpooled single-cell image galleries with assigned barcodes, cellPool facilitates interpretation and follow-up.

We applied poolVis and cellPool to profile cell-cycle regulators singly and in double perturbations. We profiled nucleus, plasma membrane, microtubule, DNA damage/repair, and mitotic markers, and demonstrated low technical variability. We identified expected phenotypes and additional candidates confirmed by follow-up. Interestingly, most double perturbations lowered γH2AX more than expected, possibly because resulting early stress may enrich for damage-resistant survivors. The epistasis hits included a recently identified synthetic lethal pair (*ATM-STAG2*),^36^ functionally coupled gene pairs (*KIF4A-SMC2, KIF4A-SMC4, KIF15-MKI67, KIF15-BUB1*, etc.), as well as novel predictions for *KIF15* interactors. These initial observations warrant further mechanistic investigation to clarify how these interactions function at the molecular level.

The current analysis has certain limitations. We did not distinguish mono- and multi-nucleated cells, the integrated nuclear intensities combined area and mean marker signal, and the epistasis analysis focused on γH2AX. Multiparametric methods considering foci number, texture, etc. could enhance statistical power. To our knowledge, we presented the first large-scale double-CRISPRi imaging data, which may provide a useful benchmark for developing such methods.

In conclusion, by combining a Cre-inducible architecture with high-quality image processing and rigorous statistical modeling of protein barcode data, poolVis and cellPool provide a robust framework for imaging-based profiling of pooled CRISPR perturbations. The platform uniquely enables double-perturbation analysis with single-cell imaging readouts, offering a scalable and extensible foundation for dissecting genetic networks and their relevance to human disease.

## METHODS

### Plasmid vector design and assembly

All plasmid backbones were cloned using the Gibson Assembly method (NEBuilder Hifi DNA Assembly Master Mix, NEB E2621). DNA fragments were obtained via PCR amplification or restriction digestion of vectors available in house or sourced from Addgene, or commercial synthesis by Twist Bioscience or Integrated DNA Technologies. The CRISPR guides (with or without barcodes) were inserted into backbone vectors by Golden Gate cloning using the Esp3I restriction enzyme (NEB, R0734) and the NEBridge Ligase Master Mix (NEB, M1100; see Supplementary Table 1a-b for the insert sequences and the vectors). Plasmids were amplified by transforming the NEB Stable Competent *E. coli* (NEB, C3040). The final plasmid sequences were verified by nanopore-based whole plasmid sequencing from Plasmidsaurus or Quintara Biosciences.

### Cell engineering and tissue culture

All cell lines were sourced from the McManus lab cryovial database or The Cell and Genome Engineering Core at UCSF. The cells were cultured at 37°C and 5% CO_2_ in tissue culture incubators and routinely tested for mycoplasma contamination.

The K562 cells were cultured in advanced RPMI 1640 (Gibco, 12633012) supplemented with 2 mM L-glutamine, 10% FBS, 100 µg/ml streptomycin, and 100 units/ml penicillin. For testing the Cas12a focused vectors, K562 cells, originally derived from a clonal line expressing dCas9-SunTag and a scFV-sfGFP-VP64 fusion (Gilbert et al., 2014), were engineered with enAsCas12a using the pRDA_174 plasmid (Addgene, 136476) and a clone was selected for high Cas12a knockout activity (engineered line borrowed from the McManus lab cryovial database).

The U2OS cells were cultured in the McCoy’s 5A (Modified) medium (Gibco, 16600082) supplemented with 10% FBS, 100 µg/ml streptomycin, and 100 units/ml penicillin.

The Hek293T cells were cultured in the Dulbecco’s Modified Eagle Medium (Gibco, 11965092) supplemented with 10% FBS, 100 µg/ml streptomycin, and 100 units/ml penicillin. For testing the Cas9 and dCas9-KRAB focused vectors, Hek293T cells were engineered with the AU5-tagged Cas protein and ERT2-Cre-ERT2 expressing constructs using the piggyBac transposon system (System Biosciences, PB210PA-1; see Supplementary Table 1a for the vectors) and the jetPRIME transfection reagent (Sartorius, 101000001). The engineered cells were selected with 1000 µg/ml hygromycin B (Gibco, 10687010).

The MCF10A cells were cultured in MEGM Mammary Epithelial Cell Growth Medium (Lonza, CC-3150) supplemented with 100 ng/ml cholera toxin. They were engineered with the AU5-tagged dCas9-KRAB, hygromycin B phosphotransferase, and ERT2-Cre-ERT2 expressing cassettes using the piggyBac transposon system and electroporation using the Lonza SE 4D Nucleofector Kit (V4XC-1032), program DS-138. They were selected with 1000 µg/ml hygromycin B (Gibco, 10687010).

### Lentivirus production, transduction, and Cre induction

Lentiviral particles were produced in the Hek293T cells by co-transfecting the pMD2.G, psPAX2, and the transfer plasmids using the jetPRIME transfection reagent (Sartorius, 101000001). Supernatant was collected between 60-72 hours after transfection, cells were removed by centrifugation, and cleared supernatant was frozen at −80°C before use. All transductions of CRISPR guide RNA carrying particles were performed at MOI < 0.1. Cre recombinase was delivered to the K562 and the U2OS cells via lentiviral transduction at MOI=1. The ERT2-Cre-ERT2 engineered Hek293T and the MCF10A cells were treated with media containing 1 µM 4-hydroxytamoxifen (Sigma, SML1666) to induce Cre recombination.

### Design of protein barcode coding sequences

A heuristic algorithm was developed to design 84 barcode coding sequences with minimal homology. Each epitope tag’s amino acid sequence was chopped into short overlapping motifs of 5-6 amino acids and the *k*-mers of lengths 8-16 bp in the coding sequences for each motif were enumerated. The complete coding sequences for each epitope tag were generated by iteratively sampling and annealing the motif-wise coding sequences while avoiding picking a coding sequence whose *k*-mer set intersects with previously sampled coding sequences. Next, for every barcode, the RNA structure stability was assessed using RNAfold for all combinations of the coding sequences of the corresponding epitope tags. Finally, the coding sequences for all barcodes were picked such that each occurrence of an epitope tag in the library is coded by a unique DNA sequence and the overall stability of the RNA structures is maximized.

### Flow cytometry

All flow cytometry data was collected using the Invitrogen™ Attune™ NxT Acoustic Focusing Cytometer. Cre recombination was induced 24 hours after transducing cells with lentivirus carrying the sgRNAs. After population-wide green fluorescence stabilized indicating effective Cre recombination, the cells were treated with media containing 2.5 µg/ml Puromycin for 60-72 hours to remove unrecombined cells. Then, flow cytometry data was collected after 21 days. Cells were detached by trypsinization (if adherent) or collected from suspension, pelleted and washed with FACS buffer (STEMCELL Technologies, 20144) twice before incubating with APC-conjugated anti-CD81 antibody (BioLegend, 349509) diluted at 1:100 in FACS buffer on ice for 20 minutes. The antibody containing solution was removed by centrifugation and the cells were resuspended in FACS buffer for flow cytometry. The resulting data was analyzed by FlowJo (10.10.0).

### Pooled library design and cloning

The targeted set of genes for the single and double perturbation screens in the MCF10A cells was curated based on expert feedback. DepMap RNAi data^39^ for the MCF10A cells was used to ensure that the majority of the targeted genes were not essential genes (Supplementary Fig. 5e). The gene targets were organized in four sets randomly. Then, four oligonucleotide pools were ordered from Twist Bioscience (see Supplementary Table 1g for the pooled sequences). Each pool consisted of 84 barcoded CRISPR guides. For every gene in the screen, top two CRISPR guide sequences were obtained from the hCRISPRi-v2.1 library.^40^ The oligonucleotide pools were amplified using the KAPA HiFi PCR Kit (Roche, 07958935001) for 16 cycles starting from 2 ng single-stranded DNA templates per 50 µl reaction. The reaction products were purified using SPRI beads (Omega Bio-tek, M1378-00), and further by 4% TBE agarose gel electrophoresis. The double-stranded oligonucleotide pools were inserted into backbone vectors by Golden Gate cloning of mixtures containing vector to insert in ratio of 1:2 using the Esp3I restriction enzyme (NEB, R0734) and the NEBridge Ligase Master Mix (NEB, M1100). After initial incubation at 37°C for 90 minutes, the reaction mixtures went through 9 cycles of 20 mins at 16°C followed by 20 mins at 37°C. Finally, the reaction was heat-inactivated at 60°C for 5 mins. Plasmid libraries were amplified by transforming the NEB Stable Competent *E. coli* (NEB, C3040) and growing LB liquid cultures containing 100 µg/ml ampicillin at 30°C for 24 hrs. Coverage was estimated to be >500x per vector construct by counting colonies from plating transformed bacteria on LB agar plates. Libraries were confirmed to be of good quality by nanopore-based whole plasmid sequencing and sequencing of individual colonies picked from the plated bacteria.

### Single perturbation and double perturbation screens

The MCF10A cells engineered to express dCas9-KRAB, hygromycin B phosphotransferase, and ERT2-Cre-ERT2 were used in the single and double-perturbation screens in Figs. 3-4. The cells were maintained in hygromycin-free media throughout the screen (Lonza, CC-3150 with 100 ng/ml cholera toxin). Before starting the screen, the cells were maintained in hygromycin-free media for two passages. Then, the cells were passaged in twenty-eight 10 cm dishes at ~2 million cells per dish (one dish per minipool). After allowing cells to adhere for 24 hours, lentiviral minipools were added to each dish at MOI < 0.1. After another 24 hours, the media in each 10 cm dish was replaced with media containing 1 µM 4-hydroxytamoxifen to trigger nuclear translocation of ERT2-Cre-ERT2. The screening time was counted starting 36 hours after adding 4-hydroxytamoxifen. This was based on the observation of dimly green fluorescent cells in the minipool cultures, indicating successful Cre induction. This timepoint was designated t=0 days.

At t=0 days, the media was changed to media without 4-hydroxytamoxifen. At t=2.5 days, the media was changed to media containing 2.5 µg/ml puromycin. Next, at t=5.5 days, the media was changed to media without puromycin. Majority of the untransduced cells died after puromycin treatment and all cultures were observed to be sparsely confluent. Then, all cultures were maintained with passaging and media changes in 10 cm dishes until, at t=16 days, they were passaged in fibronectin-coated 96-well plates (Sigma, S5171; Revvity, 6055302) at ~10,000 cells/well. At t=18 days, cells were fixed with 4% PFA (Electron Microscopy Sciences, 15714). Then, they were processed for imaging.

### Multiplexed imaging

All imaging was done using the Opera Phenix Plus high-content screening system and the Harmony 4.9 software (Revvity) using a 20X 1 NA water objective and 10% overlap between neighboring image tiles. For every tile, the different channels were imaged sequentially starting from the red fluorescent side of the spectrum to blue emission. The Hek293T images were taken in the P06-1.5H-N 6-well plates from CellVis and the U2OS and MCF10A cells were imaged after plating in the Revvity 96-well PhenoPlate (Part #6055302). In all cases, the plates were coated with fibronectin (Sigma, S5171) before seeding cells at 300,000 cells per well for 6-well plate and 10,000 cells per well in the 96-well plates. The cells were allowed to grow for 48-60 hours before fixation with 4% PFA (Electron Microscopy Sciences, 15714).

The cyclic immunofluorescence protocol was followed as described before.^41^ After fixation, cells were stained with Hoechst 33342 at 1:2000 and wheat-germ agglutinin (WGA) at 2.5 µg/ml in PBS1X at room temperature for 10 minutes. Then, first imaging cycle was done after three washes with PBS1X for 5 minutes per wash. Next, the plates were treated with 140 µl/well of acidic inactivation solution containing 3% H_2_O_2_ and 20 mM HCl in PBS1X for 6 hours. The plates were washed with 200 µl/well of PBS1X thrice and treated with 140 µl/well of basic inactivation solution containing 3% H2O2 and 20 mM NaOH in PBS1X for 1.5 hours. After washing with PBS1X thrice, the inactivation of GFP and Alexa Fluor dye conjugated to WGA was confirmed by imaging. Next, cells were permeabilized with ice-cold methanol, washed with PBS1X thrice, and incubated with Intercept (PBS) blocking buffer (LICORbio, 927-70001) at room temperature for 1 hour. For every subsequent imaging cycle, cells were incubated with fluorophore-conjugated antibody mixtures overnight at 4°C, imaged using the Opera Phenix system, and the fluorophores were inactivated using the basic solution. Three PBS1X washes were done between every change of staining or fluorophore inactivation solution.

### Preprocessing of imaging data

The images of Hek293T cells in Fig. 1 were processed by ImageJ (version 2.3.0/1.5.3f) to enhance contrast and sharpen. All other images were processed by cellPool. First, Z-slices, if available, were combined by maximum projection. Then, vignetting correction was performed using the BaSiC tool^42^ in MATLAB (R2024a) under the assumption that the vignetting pattern is identical for a tile across wells. The corrected images were stitched to create a single mosaic image per well per channel per imaging cycle. For this, an adjacency graph was created with the image tiles as nodes and edges connecting overlapping tiles based on metadata. The exact magnitude of shift was estimated by phase cross-correlation of Laplace-of-Gaussian transformed images with an upsampling factor of 10.^19^ Next, every tile is positioned relative to the well center by taking a sample of paths along the graph and adding the estimated shifts along the edges. The median of all paths is used to position every tile. This approach was applied to another channel whenever one channel failed to give a reliable positional estimate and until all channels from the imaging cycle have been used. If all channels failed to give reliable positional estimate for a tile, a linear model of all confidently placed image tiles was used to predict its position. These are pasted together to create a mosaic image for each channel in each cycle. After ensuring high quality from summary quality control reports, nuclei were segmented using a custom-trained Cellpose2 model (version 2.2.3).^43^ The features of interest were quantified using scikit-image.

### Integrating features across imaging cycles

The nuclei were tracked across cycles in three phases using the Kuhn-Munkres algorithm in each phase. First, a low-to-moderate quality tracking is performed by using orientation of nuclei as the telling feature to set up the cost function. Next, under the assumption that neighboring nuclei have similar shift from one cycle to the next, the low confidence tracking mates are identified and returned to queue for tracking again in the next phase. Finally, cells are tracked expanding out from high confidence tracking mates. We found that the transformations not accounted for by phase correlation do not contribute significantly to distorting distances with respect to local references at order of 100s of pixels but become cumulatively significant relative to nuclear size over whole well mosaics. This allowed setting up a cost function based on a pair of high confidence tracking mates to find more high confidence tracking mates in their neighborhoods using Euclidean distances after centering on the starting pair. This process continued until all existing and new set of high confidence tracking mates were used to track their neighbors. The quality of tracking is summarized as feature comparisons of the tracking mates, and unpooled images showing random samples of tracking mates per well for human-in-the-loop verification.

### Benchmarking comparison of mosaicking

For benchmarking comparison, ASHLAR was used to stitch and register the images from the barcoded U2OS samples. The mosaicked Hoechst 33342 channel from the first imaging cycle were segmented using the same custom-trained Cellpose2 model (version 2.2.3) as used for cellPool. The segmentation masks were applied to all imaging channels under the assumption of perfect pixel-to-pixel registration. Then, the barcode profiles were quantified using scikit-image in Python as before. The cycle 1 nuclear masks from cellPool and ASHLAR analyses were mapped to each other using the cellPool tracking algorithm. No mapping errors were found in 30 random samples of tracking mates from each well. Fig. 2d is a comparison of the barcode staining intensities for the set of nuclei that were tracked by cellPool across all cycles, i.e., the nuclei that washed off in later cycles were excluded. The mosaicked images from ASHLAR were unpooled by cellPool to assess the mosaicking quality.

### Mixture model for debarcoding protein barcodes

The median intensities taken over a nuclear mask give the barcode profile denoted 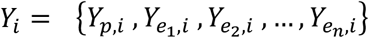 for the corresponding cell *i*, where *p, e*_1_, …, *e*_*n*_ are the fluorescent barcode backbone and the *n* epitopes in the library, respectively. *Y* represents the binary logarithm of median intensity. Let 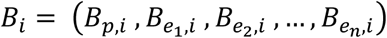 denote a barcode, where *B*_*e,i*_ is a state variable. *B*_*e,i*_ = 1 when cell *i* is positive for the epitope tag *e* and 0 if it is negative. cellPool takes the barcode profiles as input and outputs the barcodes in two stages with the first being optional.

#### Population-level thresholding

This step deciphers the barcodes completely for some cells and partially for the rest by analyzing the intensity distribution for each epitope across the population. By default, we apply Li’s iterative method based on minimum cross-entropy^44^ to determine a threshold, *t*_*p*_ for GFP signal. We classify *B*_*p,i*_ as 1 if *Y*_*p,i*_ is greater than *t*_*p*_. By default, if *Y*_*p,i*_ is less than *t*_*p*_ − 1, we classify *B*_*i*_ = (0, 0, …, 0). The default settings can be changed in the cellPool configuration profile to values determined by manually studying the histograms generated by cellPool. The thresholding method can be set to any method from the *filters* submodule of scikit-image. We assume that enough GFP-negative cells (order of 1000s) are available collectively on the plate. If not, cellPool skips to multivariate modeling directly.

For the GFP-positive cells, we determine high-confidence thresholds for each epitope by comparing the epitope intensity distribution in the GFP-positive cells with that in the GFP-negative cells (Fig. 2h). Say, 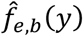 is the empirical density function of the background noise for epitope tag *e* as quantified using the GFP-negative cells. Let *w*_*e*_ be the proportion of GFP-positive cells that are also positive for the epitope tag *e*, and *f*_*e,s*_(*y*) be the probability density function for the log-transformed staining intensity of *e* in these cells. We determine the high-confidence threshold *t*_*e*_, such that for 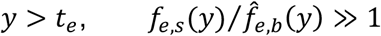. This happens when 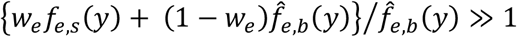. Here, 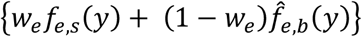 is the signal-background mixture in the GFP-positive cells. Hence, cellPool estimates 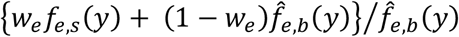 for each epitope tag and positions the threshold such that the ratio is consistently greater than a user-specified level.

#### Multivariate mixture model-based debarcoding

To account for distributions with non-elliptical isodensity loci, we partition the GFP-positive cells into 5-10 sets that have relatively uniform GFP intensity and thereby, low variation in epitope intensities as well. This permits a good fit by a mixture model based on skew elliptical distributions to the data for each set. We model the single cell intensity vectors as a mixture distribution with as many components as there are barcodes. Let ***e***_***k***_ be an *n* × 1 one-hot vector representing the epitope *e*_*k*_, *m* be the number of epitopes per barcode, 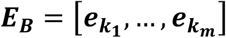 be an *n* × *m* matrix with one-hot vectors for epitope tags in the barcode *B* as its columns, and 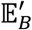 be the set of absent epitopes, i.e., 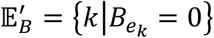. Then, for barcode *B* and 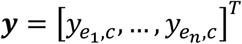, we model the observed distribution as

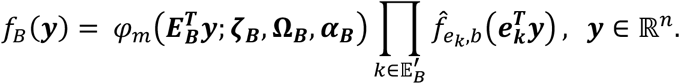

Here, *φ*_*m*_ is the *m*-dimensional multivariate skew normal distribution with **ζ**_***B***_ ∈ ℝ^*m*^, **Ω**_***B***_ ∈ ℝ^*m*×*m*^, and ***α***_***B***_ ∈ ℝ^*m*^ as the location, scale and skewness parameters, respectively. This equation describes a multivariate skew normal distribution in the subspace of epitope tags present in barcode *B* and empirically estimated background distribution in the subspace of the remaining epitope tags. cellPool fits a mixture model with components defined by the above equation (Supplementary Fig. 3). It uses the expectation-maximization algorithm with the option to test multiple parameter initializations in parallel and quality control the convergence of the iterations of optimization to maximize the data likelihood,

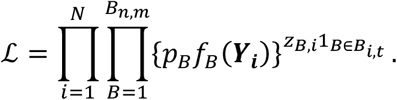

Here, *N* is the total number of nuclei, *B*_*n,m*_ is the total number of barcodes, *p*, is the mixing fraction of barcode *B, z*_*Bi*_ is an indicator variable that is 1 if *i* has barcode *B* and is 0 otherwise, and 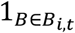 is an indicator variable that equals 1 if *B* is compatible with the epitope set *B*_*i,t*_ present in *i* based on population-level thresholding and 0 otherwise.

cellPool initializes the model parameters as follows. The skewness parameters are set to random values, off-diagonal terms in the scale parameters are set to 0, the location parameters are set to the foreground mean based on Otsu’s thresholding, and the mixing proportions are set as equal for all barcodes. Then, cellPool iteratively computes the conditional expectations for the latent variables (E-step) and updates the parameters for the multivariate skew normal components (M-step) following the equations described before,^20^ except that the conditional expectations for *z*_*Bi*_ and likelihood calculations use our model for protein barcode data (see *f*_*B*_(***y***) and *ℒ* above). The EM algorithm terminates upon convergence of log-likelihood and outputs the best parameter estimates, which yield the posterior probabilities for all possible barcode assignments. cellPool assigns a barcode if the corresponding posterior probability meets the user-defined threshold.

### Benchmarking comparison with other debarcoding methods

cellPool’s debarcoding model was benchmarked by comparison with the Single-Cell Deconvolution (SCD)^21^ and the EPICode^6^ methods using custom scripts. For SCD, each epitope tag’s raw signal was normalized with respect to the 99^th^ percentile of its distribution encompassing all cells under consideration. Then, the top 3 epitope tags present in a cell were assigned as its barcode if the 3^rd^ and 4^th^ highest ranked epitope tags met the *barcode separation* criteria, i.e., their difference was greater than a user-defined threshold. For EPICode, each epitope tag’s raw signal was converted into a percentile score based on the background distribution. Then, for each barcode, a score was calculated as the mean percentile score of its constituent epitope tags. The barcode with the highest score was assigned if the difference from the next highest score meets the barcode separation criteria. For cellPool, the difference in the posterior probabilities assigned to the top two barcodes was used to quantify confidence. The barcode separation criteria was varied from 0 to 1 to generate the curves in Fig. 2k.

### Validation of guides for double perturbation screen

Top ranked guides for *ATM, KIF15, KIF22*, and *KIF4A* were validated in the MCF10A cells engineered with dCas9-KRAB and ERT2-Cre-ERT2. The targeting guides were inserted under the hU6 promoter and the nontargeting under the mU6 promoter in the same poolVis vector. After 18 days of screen, the cells were fixed in 96-well plates as described before and imaged in two cycles. In the first cycle, Hoechst 33342 and GFP signals were captured. Then, GFP was denatured using protocol described for cyclic immunofluorescence above. Then, the samples were permeabilized using ice-cold methanol and incubated with Intercept (PBS) blocking buffer (LICORbio, 927-70001) at room temperature for 1 hour. Then, the samples were incubated with primary antibodies for 24 hours at 4°C, followed by another round of incubation with secondary antibodies for 24 hours at 4°C. The samples were washed, stained with Hoechst 33342 and imaged again.

*ATM, KIF22*, and *KIF4A* are nuclear localized during interphase while *KIF15* is cytosolic. Additionally, *KIF22* and *KIF4A* are expressed primarily during the late G2/M phase. *ATM* signal distribution was quantified by the median nuclear staining intensities of its antibody. *KIF22* and *KIF4A* signal distribution was quantified as median nuclear staining intensities after thresholding for late G2/M phase nuclei using the Hoechst 33342 signal. *KIF15* signal distribution was quantified as median cytosolic staining intensities after obtaining the cytosolic mask as a strip of width 3 pixels (1.8 µm) encircling the nuclear mask.

### Phenotypic feature extraction and differential analysis

Phenotypic features were extracted using scikit-image in Python. The nuclear area was computed for objects identified by Cellpose2 segmentation of images. The integrated nuclear γH2AX and H3S10P signal was obtained as the sum of fluorescent intensities over nuclear masks. Mean cytosolic α-tubulin signal was obtained as the mean fluorescence intensity in the corresponding image channel and the cytosolic mask represented by a strip of width 3 pixels (1.8 µm) encircling the nuclear mask. For single perturbation screen, log10-transformed intensities were used in differential analysis. For double perturbations, the log10-transformed intensities were further centered and scaled with respect to the median and median absolute deviation of the untransduced GFP-negative cells for every column of every imaged plate. Differential analysis was performed using the Student’s t-test to compare normalized feature for a target group vs the nontargeting control infected cells. The p-values were corrected for multiple testing by the Benjamini-Hochberg procedure. The effect of nontargeting controls was centered at 0 and standard errors were used to scale the mean effect sizes of perturbations.

For mitotic index analysis, the mitotic nuclei were identified by manually thresholding integrated nuclear γH2AX. Then, the mitotic index was measured for each targeting sgRNA as the proportion of mitotic nuclei relative to all nuclei identified with that target. The mitotic indices for well-covered guides (coverage > 1500) were compared with the mitotic index of the combined non-targeting and untransduced control population by two-sided exact binomial test. The p-values were adjusted for multiple testing using the Benjamini-Hochberg procedure. The hits were identified as the gene targets with FDR <0.1 and same sign of effect for both guides.

Genetic interaction scores were quantified as difference of double knockdown effect and the sum of single knockdown effects.^30^ Interaction hits for DNA damage and repair were identified as those gene pairs where the estimates for single knockdown effects were positive for both guides targeting the Cre inducible gene perturbation and the double knockdown effect was negative with FDR-adjusted p-values <0.2. The representative images in the figures were unpooled by cellPool.

## DATA AVAILABILITY STATEMENT

### Data and reagent availability

The datasets generated during this study will be deposited in AWS and made publicly available upon acceptance. Plasmid constructs will be deposited on Addgene and made publicly available upon acceptance. Any additional materials or information related to the study can be obtained by reaching out to the corresponding authors with a reasonable request.

### Code availability

All code used in this article is available at:https://github.com/mcmanuslab/cellpool and https://doi.org/10.5281/zenodo.17577408.

## ACKNOWLEDGEMENTS

We thank members of the McManus lab for helpful discussions, technical support, and feedback throughout this project. This work was assisted by the UCSF Parnassus Center for Advanced Technologies. A special thanks to Iain Cheeseman, Monica Bettencourt-Dias, Mariana Lince-Faria, Lee Rao, Shalin Mehta, Talon Chandler, Eduardo Hirata-Miyasaki, Soorya Pradeep for many helpful discussions. This study was supported, in part, by National Institutes of Health grants U01CA272546 and U24DK116214 to M.T.M., as well as a Sandler Program for Breakthrough Biomedical Research Award to M.T.M. Additional support was provided by the Chan Zuckerberg Biohub, Pivotal Life Sciences, and the Diabetes Center at UCSF to M.T.M.

## AUTHOR CONTRIBUTIONS

K.C. and M.T.M conceived the project. K.C. and M.T.M designed the wet-lab method prototype. K.C. developed the wet-lab methods with critical input from M.T.M. K.C. developed the computational methods and the software. K.C. performed the experiments, assay optimizations, and analyzed data with feedback from M.T.M. K.C. drafted the original manuscript. K.C. and M.T.M. edited the manuscript. Both authors reviewed and approved the final manuscript.

