## Supplementary Figures 1-9, Supplementary Tables 1-4, Supplementary Videos 1-5, and Additional Data 1-2 for "Scalable imaging-based profiling of CRISPR perturbations with protein barcodes": Supplementary_Information.pdf

### LIST OF SUPPLEMENTARY FIGURES

1. Flow cytometry data to develop the poolVis vectors and the TATAlox versions of the human U6 promoter
2. cellpool quality control measures show high-quality image mosaicking
3. Example of a quality control summary for image mosaicking by cellPool
4. Plate diagram for the cellpool debarcoding model
5. Characterization of the dCas9-KRAB engineered MCF10A cells and other single-perturbation screen data
6. Validation of single-perturbation screen by comparison with a screen of arrayed individual guides
7. Validation of guides for *ATM* and the chromokinesins in the dCas9-KRAB engineered MCF10A cells
8. Normalization and quality control of the features from the double perturbation screen
9. Additional data from the double perturbation screen

### DESCRIPTIONS OF SUPPLEMENTARY VIDEOS

Supplementary Video 1. Stitching and registration performance of ASHLAR as characterized by applying the nuclear segmentation masks from cycle 1 to the later cycles. Each frame in the video is showing a random sample of 30 masks from the well annotated at the bottom of the frame. Each square in the grid layout has 90 µm long sides.

Supplementary Videos 2 and 4. Quality control of image mosaics generated by cellPool. Each frame is showing 90 µm wide strips centered on the edges of the image tiles with the three highest standard errors of mean positional estimates in a well. Poor quality stitching, if any, will show as misaligned nuclei at the center of the displayed strips. Supplementary Video 2 is from cellPool analysis of the U2OS data in Figure 2. Supplementary Video 4 is from cellPool analysis of the MCF10A data in Figure 3.

Supplementary Videos 3 and 5. Quality control of tracking performance of cellPool. Each frame is showing a random sample of 30 tracked nuclei from the well annotated at the bottom of the frame. Each square in the grid layout has 180 µm long sides. Supplementary Video 3 is from cellPool analysis of the U2OS data in Figure 2. Supplementary Video 5 is from cellPool analysis of the MCF10A data in Figure 3.

### DESCRIPTIONS OF SUPPLEMENTARY TABLES

Supplementary Table 1. Descriptions of tables listed in Sheet 1 of the Excel Sheet.
Supplementary Table 2. Results of differential analysis for the single perturbation screen.
Supplementary Table 3. STRINGDB data for subset of *KIF15* interactors.
Supplementary Table 4. Results of differential analysis for the double perturbation screen.

### DESCRIPTION OF ADDITIONAL DATA 1

A collection of images unpooled by cellPool for the interaction of *KIF4A* and *RACGAP1*. The image file names indicate the pair of perturbations represented in the images and the phenotypic classification. Each square panel in the files is 120 µm x 120 µm. The magenta color is showing the plasma membrane stain (WGA) and the green color is showing the nucleus (Hoechst 33342). The cell/nucleus at the center of each square panel is associated with the knockdown targets in the file name. Other neighboring cells may have other perturbations, not labeled in these images. In multinucleated cells, the nuclei in the cell at the center were debarcoded individually and confirmed to have the same barcodes.

### DESCRIPTION OF ADDITIONAL DATA 2

Vector maps for the plasmid vectors developed in this manuscript. See Supplementary Table 1 for plasmid descriptions.

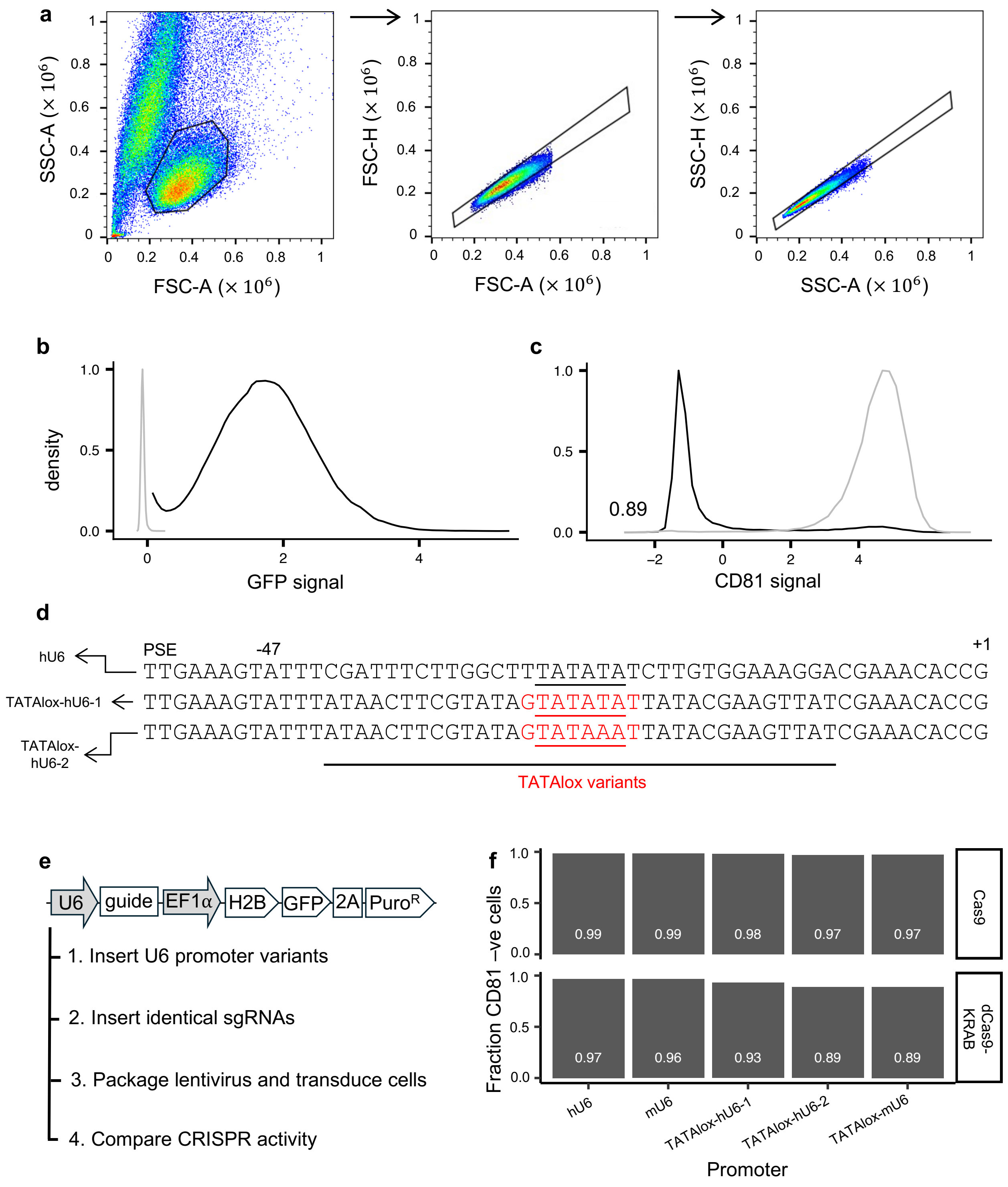


**Supplementary Figure 1. Flow cytometry data to develop the poolVis vectors and the TATAlox versions of the human U6 promoter. (a)** Example of the typical gating strategy for the flow cytometry data. Single cells are identified by gating on the SSC vs FSC scatterplot followed by gating on the FSC-H vs FSC-A and SSC-H vs SSC-A. **(b)** GFP signal from the design in Figure 1e, scaled and centered with respect to the background green fluorescence in untransduced control Hek293T cells (gray). **(c)** *CD81*knockdown signal from the design positioned in Figure 1e, scaled and centered with respect to the non-targeting control in the same Hek293T cells engineered with dCas9-KRAB (gray). **(d)** TATAlox U6 variants showing the lox sites (underlined) with the TATA motif embedded in its spacer region (red text) and positioned downstream of the proximal sequence element (PSE) of the U6 promoter. **(e)** Vector backbone and experiment design for testing the TATAlox-U6 promoter variants. **(f)** Bar plots comparing the effectiveness of the promoter variants in context of wild-type Cas9- and dCas9-KRAB-based *CD81* knockout and knockdown in the Hek293T cells, respectively.

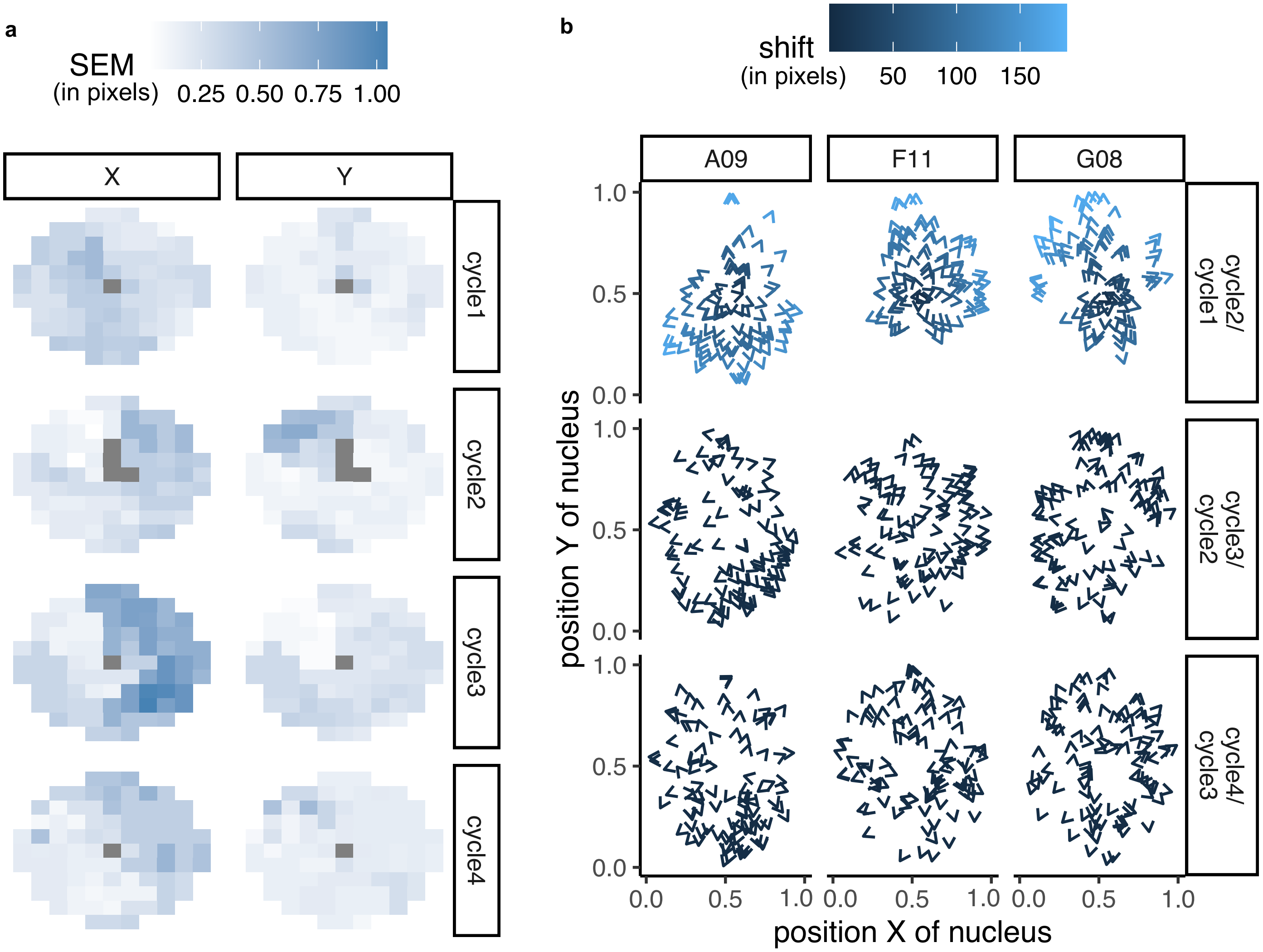


**Supplementary Figure 2. cellpool quality control measures show high-quality image mosaicking. (a)** The colormap gives the standard error of mean (SEM) in the estimated position of imaging tiles. Each circular tiling pattern is showing the SEM (in pixels) in the X- or Y-coordinates for every tile in the mosaic image per well per cycle. The plot shows an example well mosaicked by cellpool. The central tile is always centered at the center of the mosaicked image. The grey tiles indicate unavailable SEM values due to a single positional estimate being available. **(b)** The shift (in pixels) between cellpool tracking mates observed for randomly downsampled nuclei from three randomly selected wells.

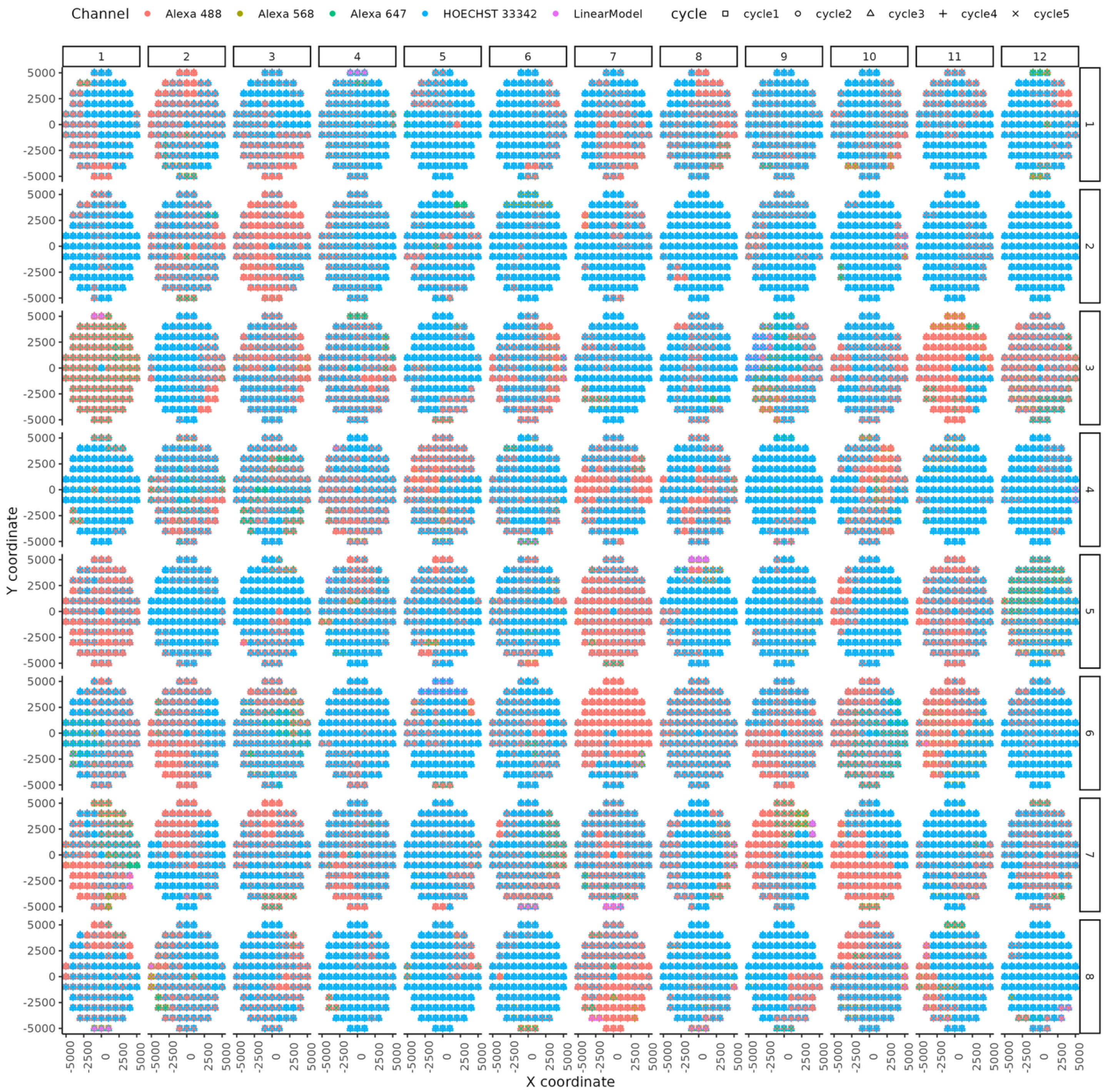


**Supplementary Figure 3. Example of a quality control summary for image mosaicking by cellPool.** Each point is showing the positional estimate for an image tile in a cycle and the imaging channel that provided the estimate. Poor quality alignment appears as unaligned shapes for the same tile or excessive use of predicted positions based on a linear model. Typical reports show frequent use of a mix of imaging channels to accurately position all tiles.

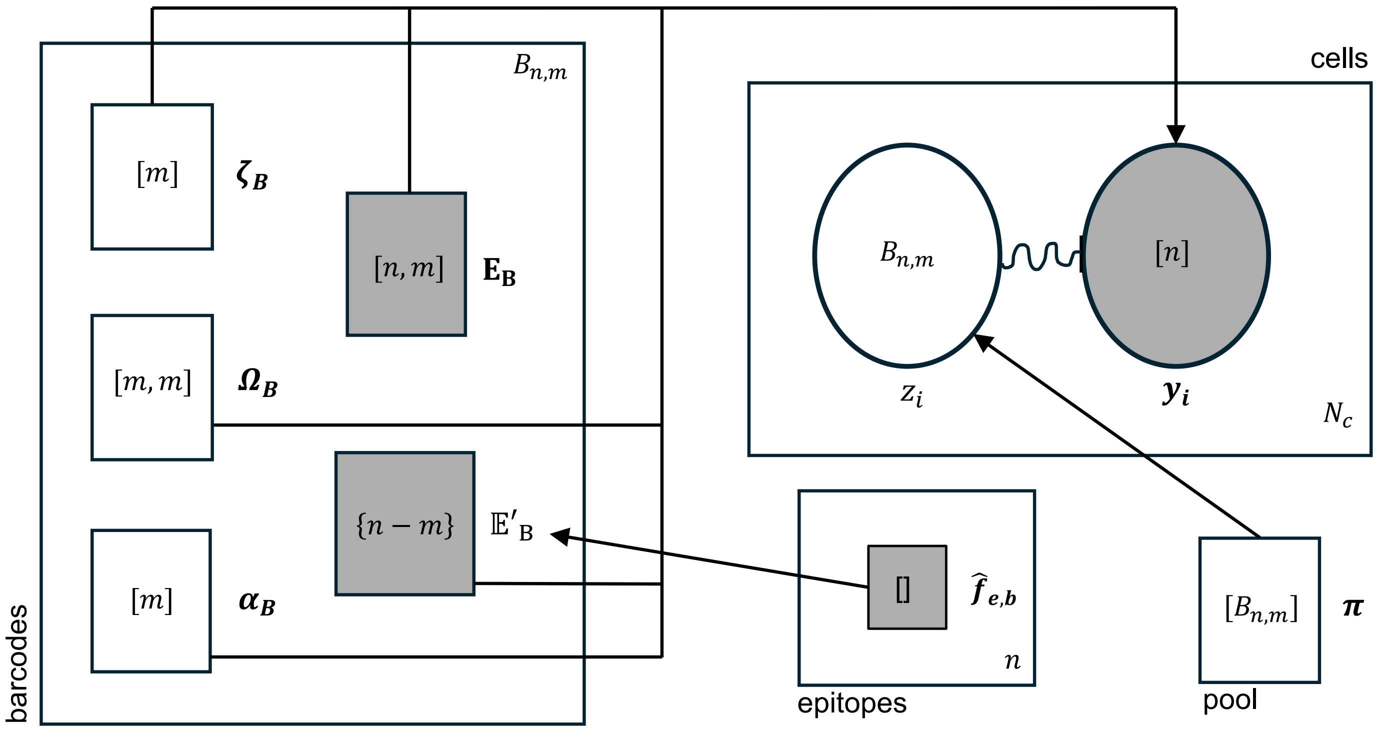


**Supplementary Figure 4. Plate diagram of cellpool's probabilistic generative model for the barcode profiles.** The cellpool debarcoding model consists of parameters associated with four experimental components relevant to the barcode data– the pooled sample, single cells, protein barcodes, and the epitopes. The large plates labeled with each component contain the parameters associated with that component, shaded objects represent known variables, and the arrows show dependencies between the parameters and observations. The quantity at the top/bottom right corner of a plate represents the number of variations for that component, e.g., $n$ epitopes total in the library, $B_{n,m}$ barcodes by choosing $m$ epitopes per barcode, and $N_c$ cells observed in the experiment. From the perspective of deciphering the protein barcodes, all wells are considered as a single pool.

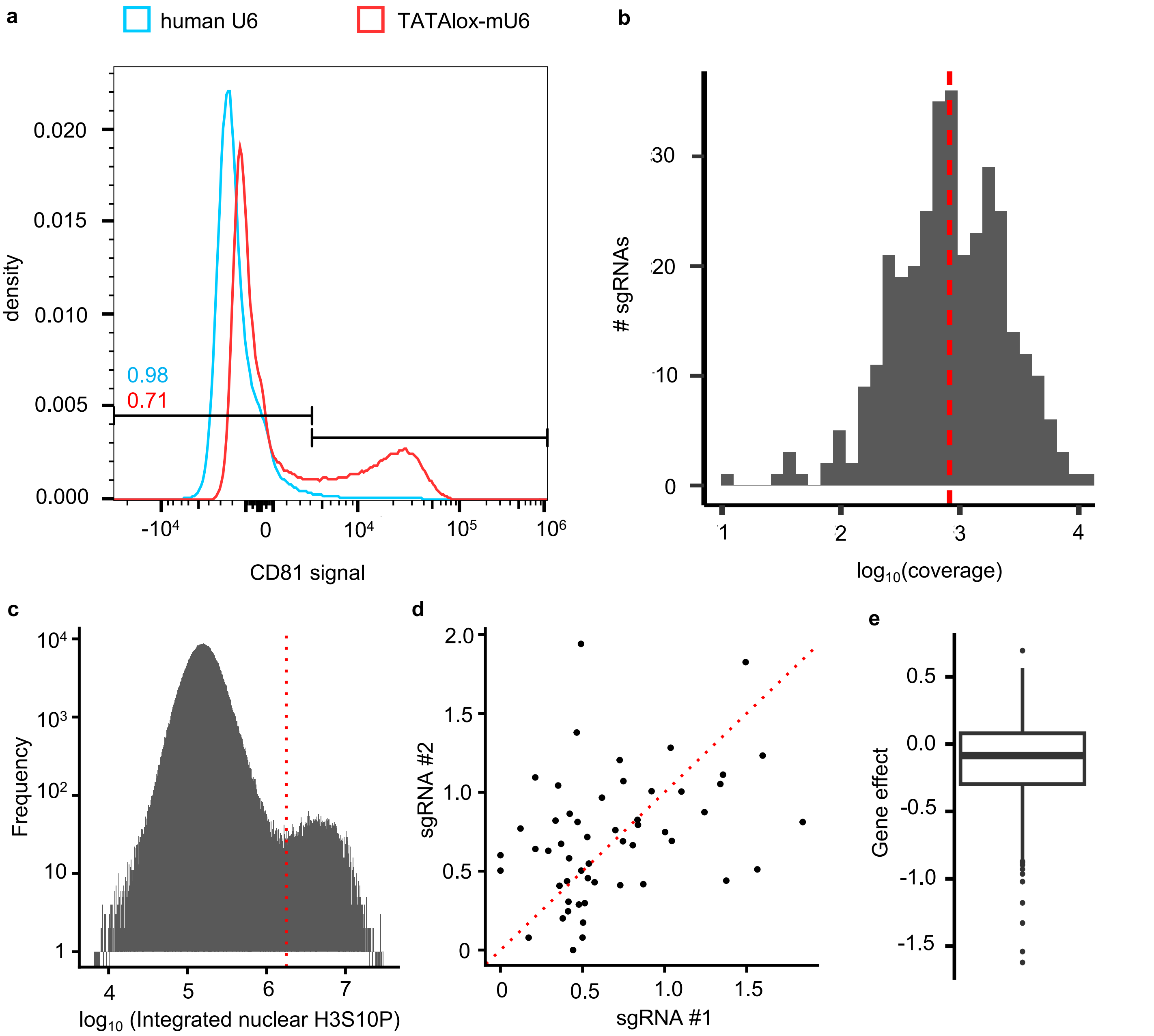


**Supplementary Figure 5. Characterization of the dCas9-KRAB engineered MCF10A cells and other single-perturbation screen data. (a)** Distribution of *CD81* signal (biexponential transformation) when the sgRNA for *CD81* knockdown is driven by the human U6 promoter in the MCF10A cells vs the TATAlox mouse U6 promoter. **(b)** Coverage distribution for sgRNAs in the single perturbation screen. Red dashed line is marking the median coverage. **(c)** Histogram showing the distribution of integrated nuclear H3S10P signal. Dotted red line is showing the manually determined threshold for mitotic cells. **(d)** Scatterplot showing the mitotic indices measured from the two guides for each of the 53 genes with mean coverage greater than 1500. **(e)** Boxplot showing the distribution of effects of the targeted gene knockdowns from the DepMap RNAi data in the MCF10A cells. Scores less than -0.5 represent depletion and scores less than -1 represent strong lethality.

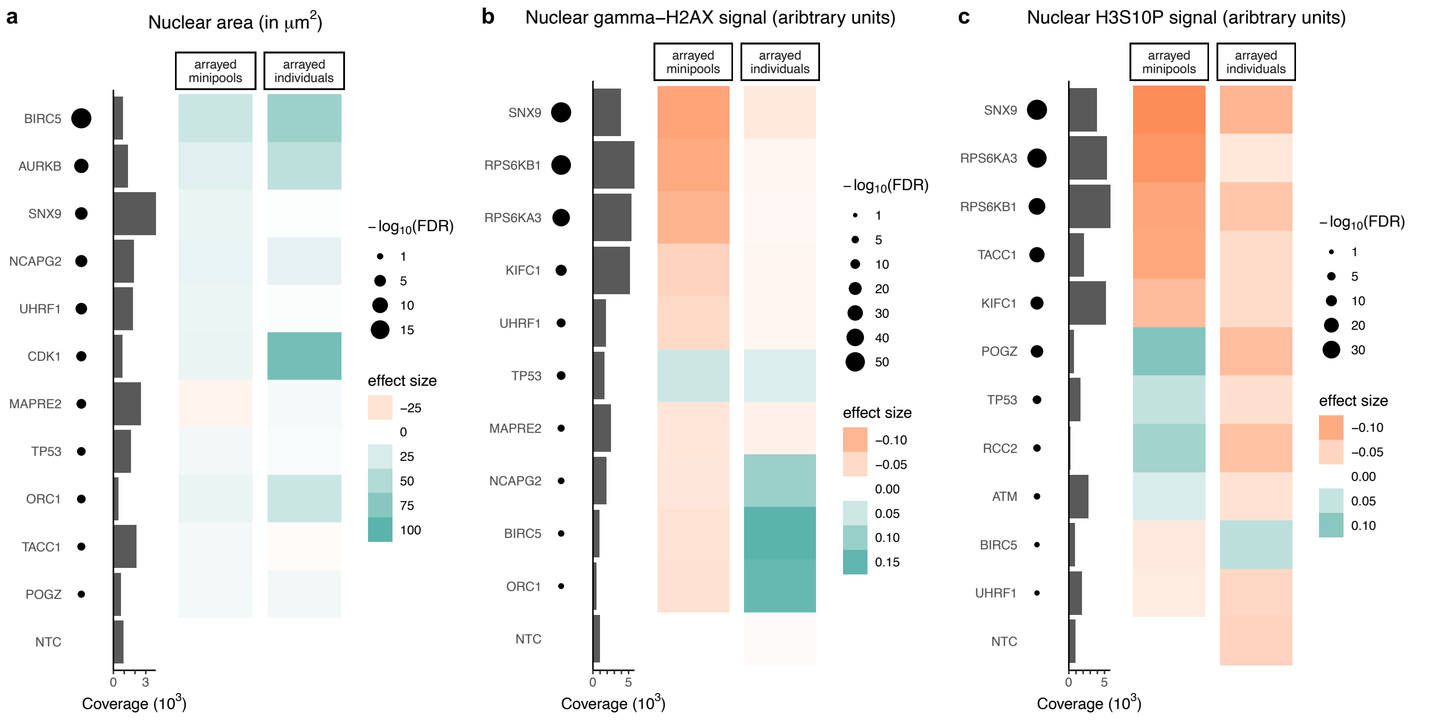


**Supplementary Figure 6. Validation of single-perturbation screen by comparison with a screen of arrayed individual guides. (a-c)** Heatmaps showing comparison of estimated effect sizes for various perturbations in the arrayed minipools screen vs the follow-up screen of arrayed individual guides. Each row shows one gene target. The nontargeting control (NTC) is shown in the bottom row. The annotations with black circles and barplots give the main quantities determining confidence in the arrayed minipool estimate for each target– the FDR-adjusted *p*-values and the coverage. When the coverage and FDR-adjusted p-values are both high, the arrayed minipools-based estimates of knockdown effect on the features in panel titles are overall consistent with the estimates from the screen of fully arrayed individual sgRNAs.

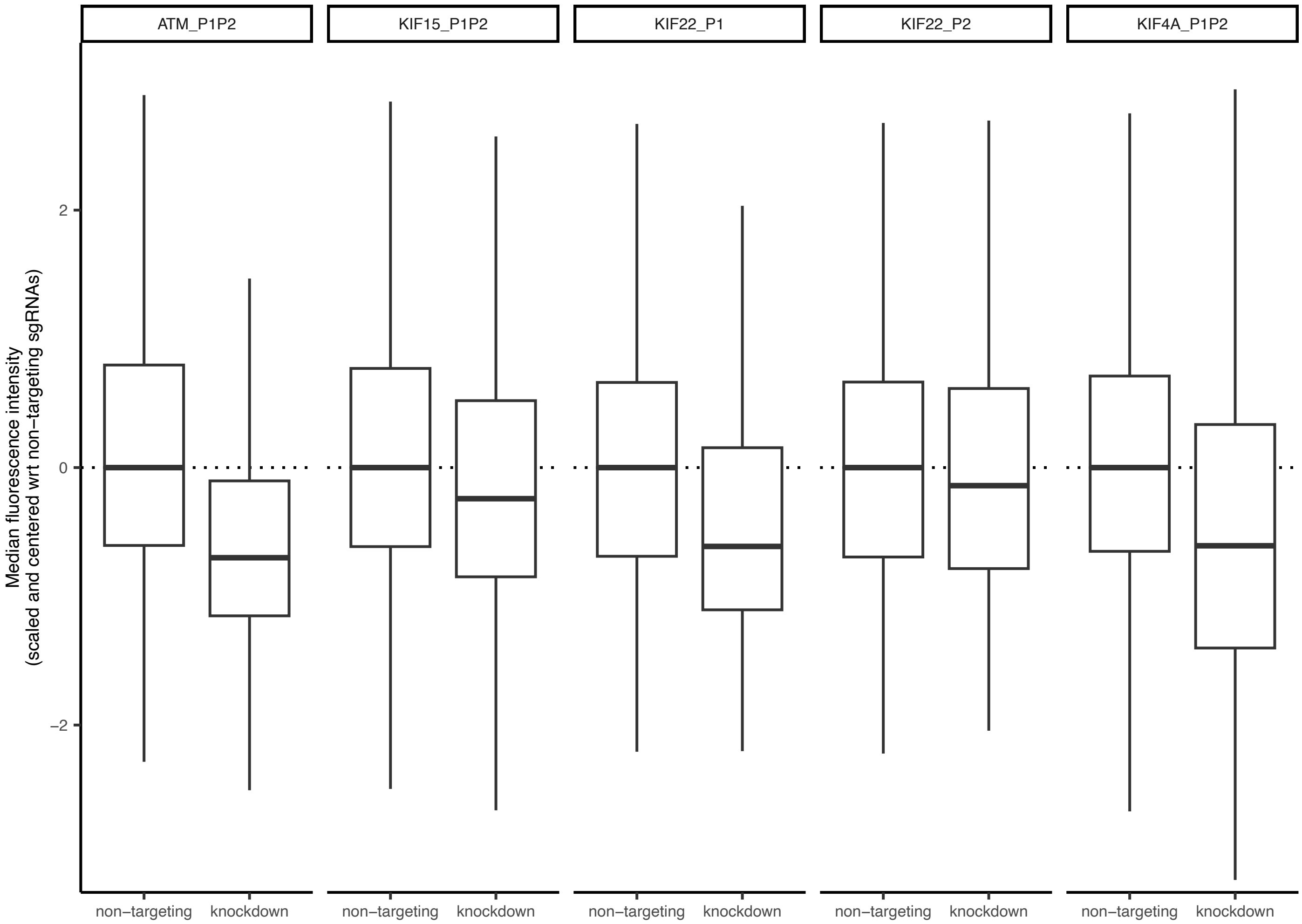


**Supplementary Figure 7. Validation of CRISPR guides for *ATM* and the chromokinesins in the dCas9-KRAB engineered MCF10A cells.** Boxplots showing knockdowns in fluorescence intensity distributions from the gene targeting guides.

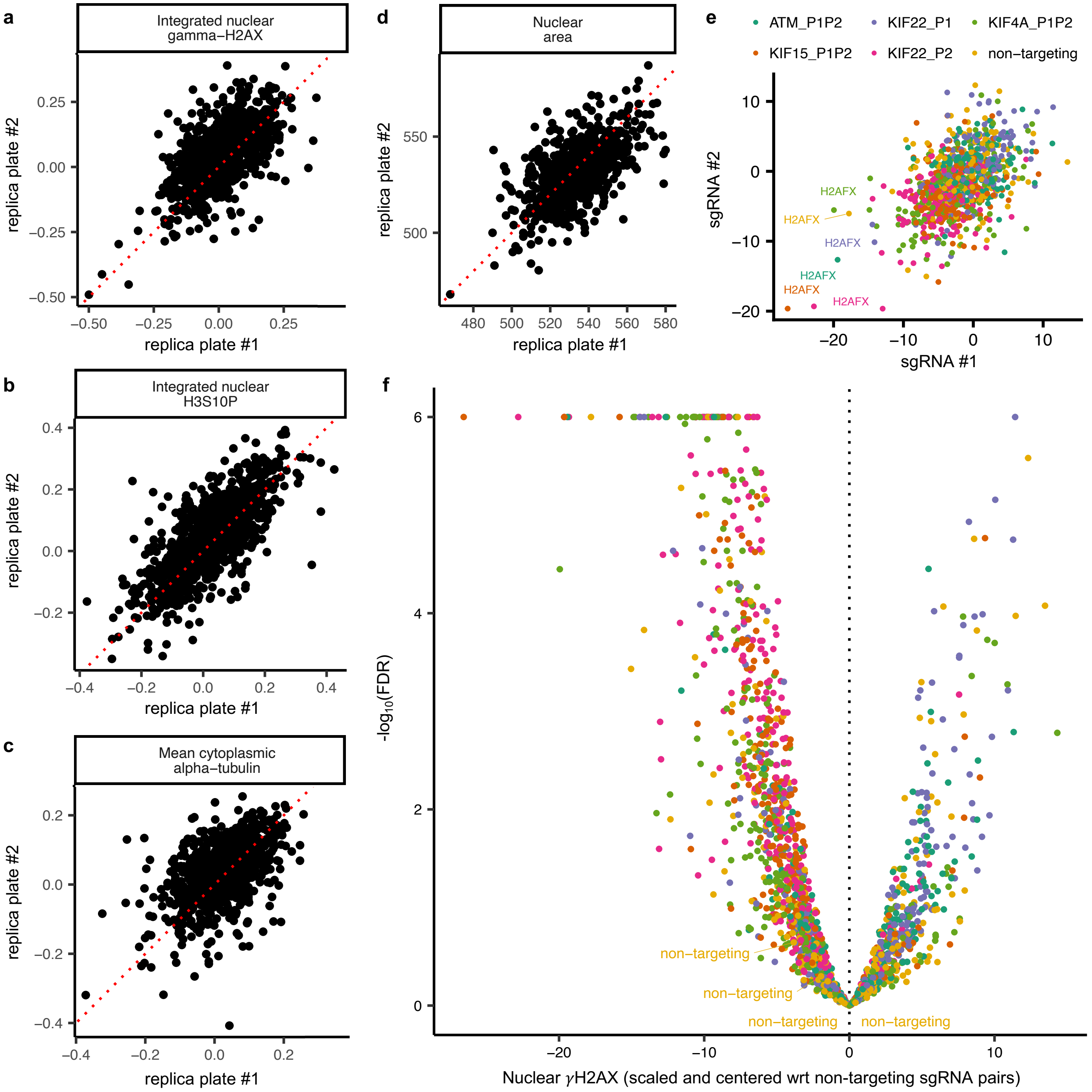


**Supplementary Figure 8. Normalization and quality control of the features from the double perturbation screen. (a-d)** Comparison of mean normalized feature value for the CRISPR perturbation combinations in the two replicate plates for each sample. The red dotted lines are the identity lines. **(e)** Comparison of knockdown effect on nuclear $\gamma$H2AX signal from the two guides targeting the same gene in our *guide2* minipools. Colors represent the *guide1* targets. **(f)** Volcano plot for the nuclear $\gamma$H2AX signal. Color mappings are identical to panel e.

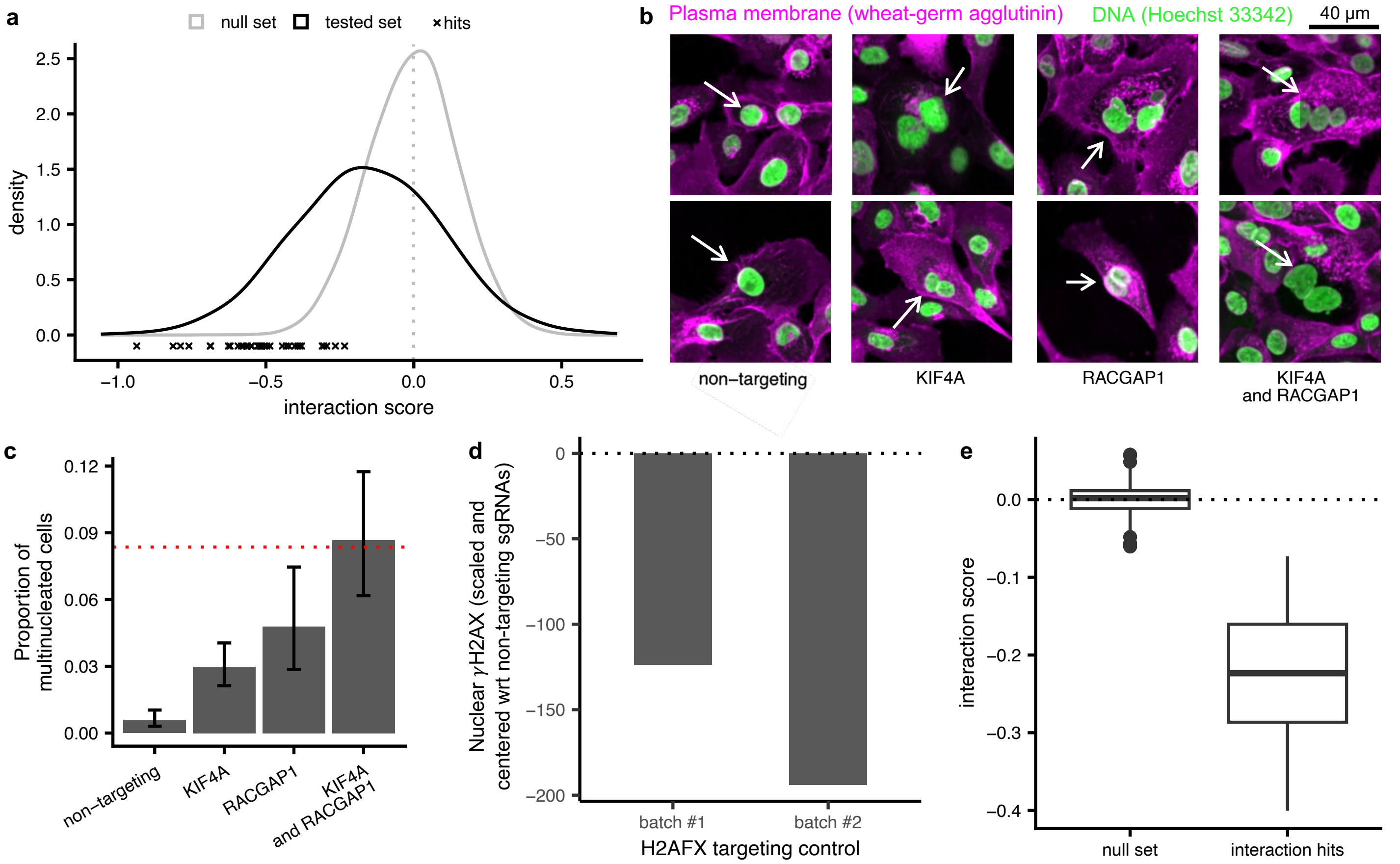


**Supplementary Figure 9. Validation of the genetic interaction hits by comparison with a fully arrayed interaction screen. (a)** Distribution of genetic interaction scores (normalized with respect to the absolute magnitude of γH2AX knockdown effect size) for the null set from bootstrapping non-targeting control cells (gray) and the tested perturbation combinations. The points shaped "x" close to the x-axis represent the selected high-confidence hits based on reproducibility of interaction pattern across both guides targeting the same gene. **(b)** Representative images showing cytokinesis defects. **(c)** Bar chart showing the proportion of multinucleated cells. The error bars are marking the 95 % binomial confidence intervals. **(d)** Robust knockdown of nuclear γH2AX signal is observed in both batches of H2AFX targeting controls alongside the arrayed validation experiment. **(e)** Boxplot showing the distribution of interaction scores (normalized with respect to the absolute magnitude of γH2AX knockdown effect size) for the null set from bootstrapping non-targeting control cells and the fully arrayed interaction screen.
