## Supplementary figures and images for "Scalable imaging-based profiling of CRISPR perturbations with protein barcodes"

### KIF4A_nontargeting_multinucleated.png

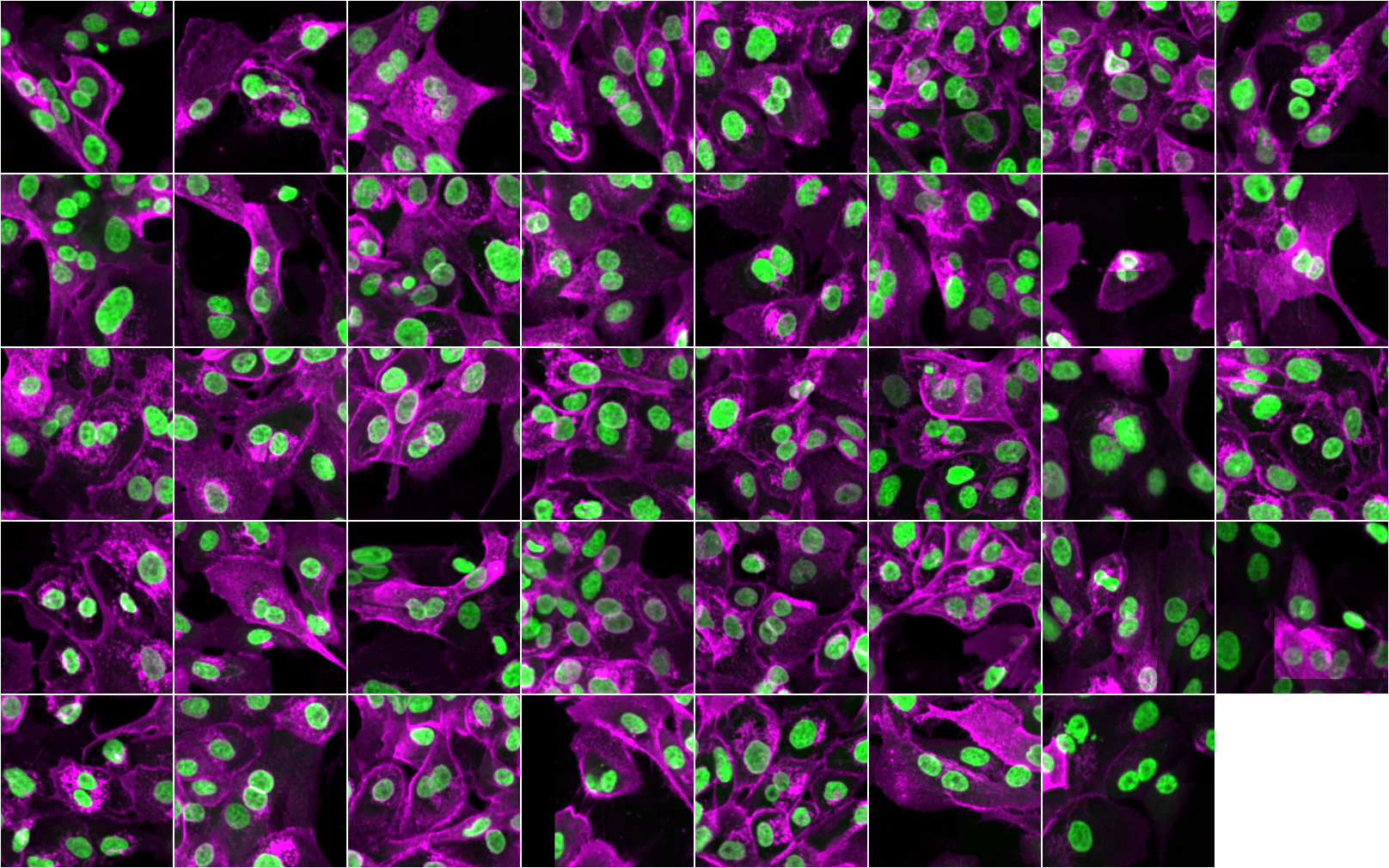

### KIF4A_RACGAP1_multinucleated.png

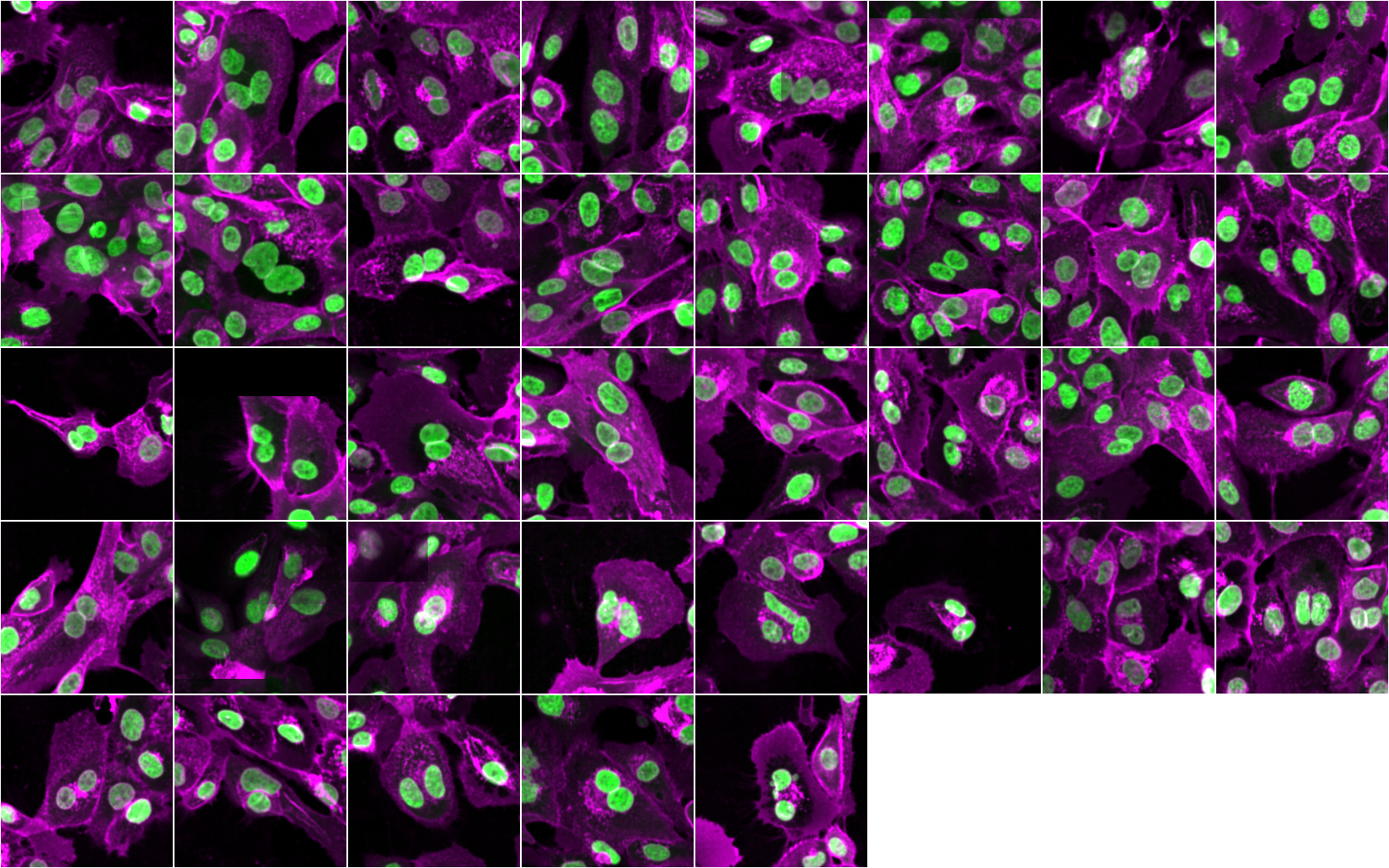

### KIF4A_RACGAP1_regular.png

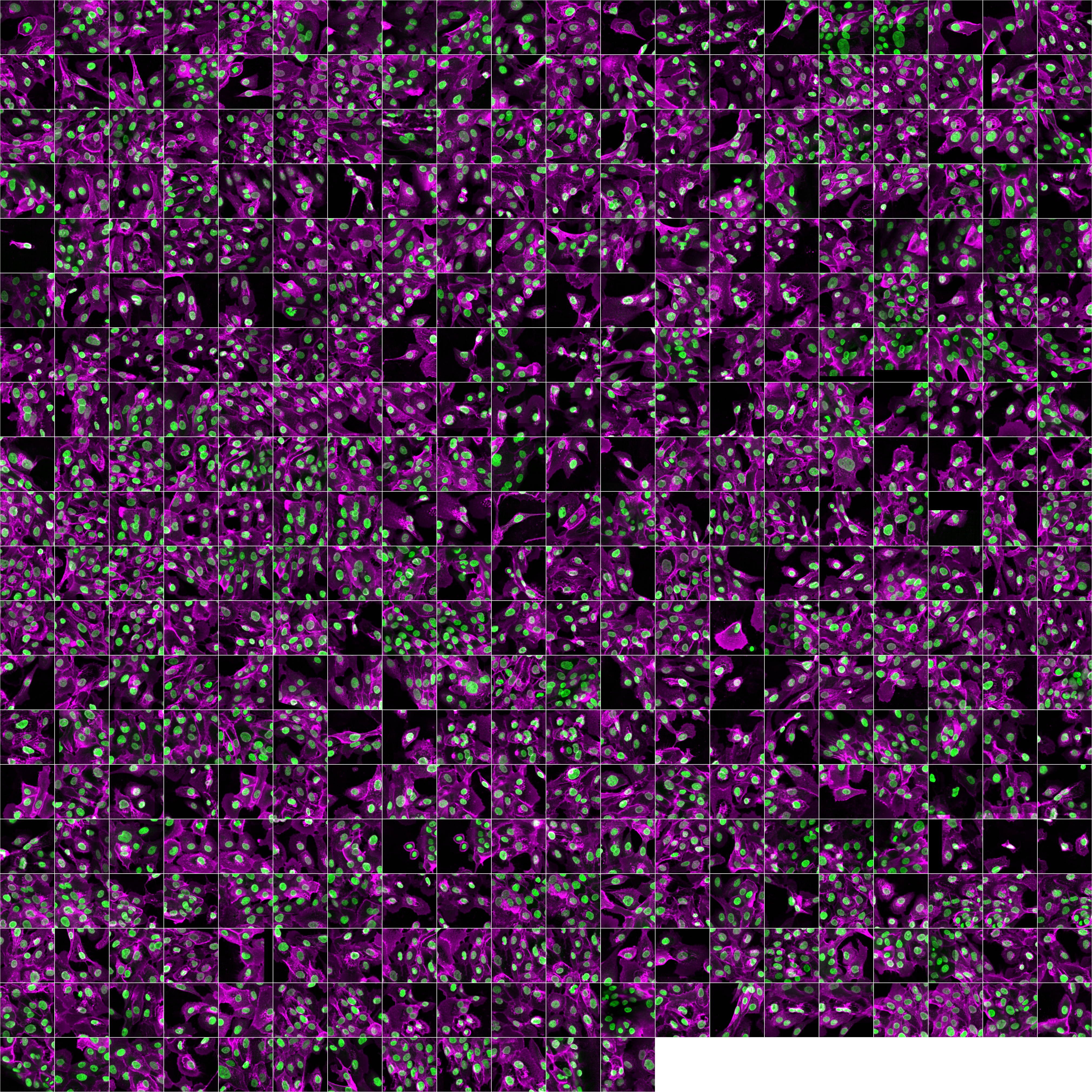

### NTC_1__nontargeting_multinucleated.png

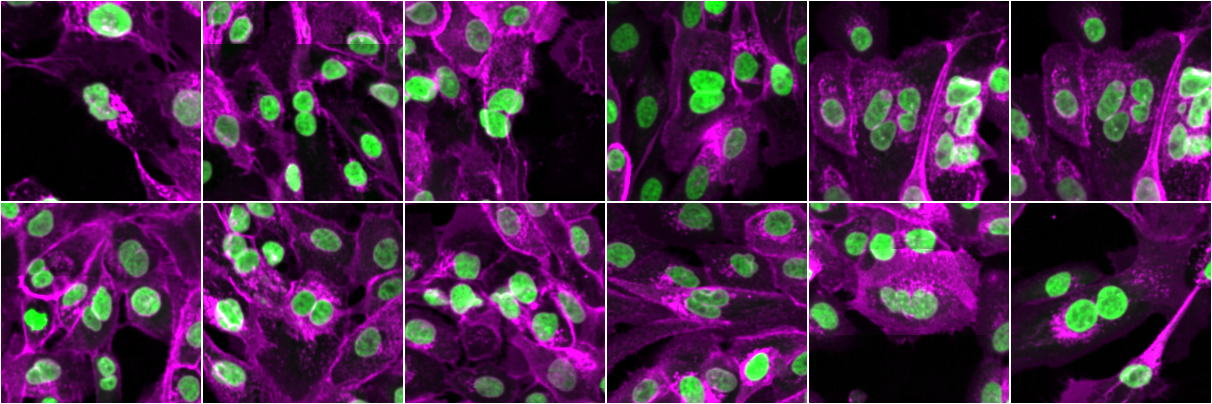

### NTC_RACGAP1_multinucleated.png

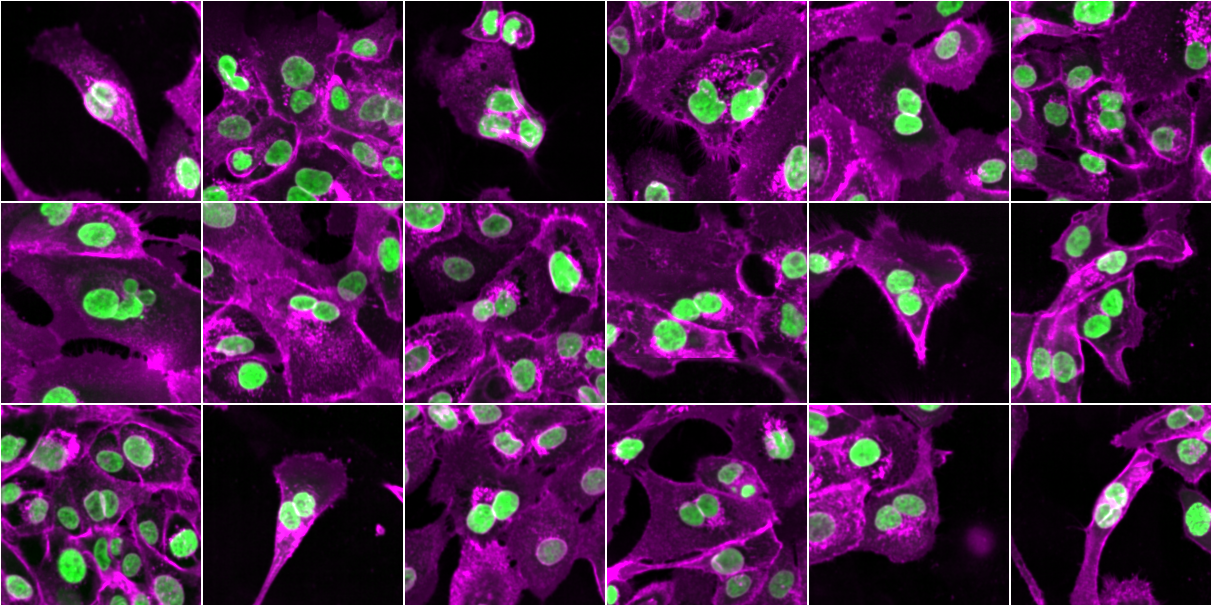

### NTC_RACGAP1_regular.png

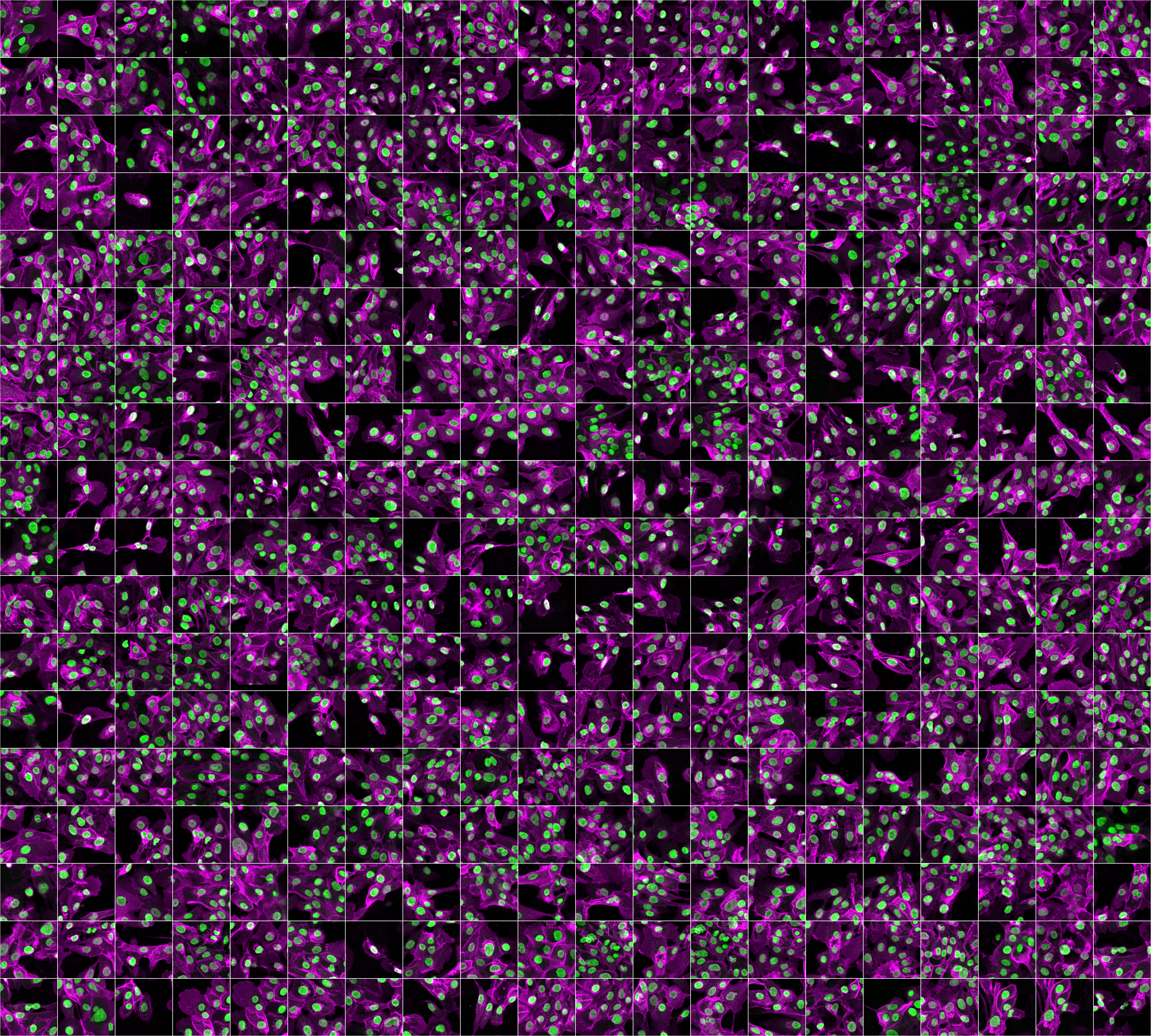
